# Interspecific social contacts spread gut microbiota in wild rodent communities

**DOI:** 10.64898/2026.11.04.764353

**Authors:** Aura Raulo, Eveliina Hanski, Curt Lamberth, Josh A Firth, Sarah CL Knowles

## Abstract

**Background:** The mammalian gut microbiota influences a wide array of host phenotypes and varies substantially among individuals. Among conspecifics, microbiota composition is strongly shaped by microbial transmission through social interactions, yet the extent to which such transmission also occurs across species boundaries remains poorly understood. Here, we combine high-resolution tracking data collected over more than 10 months with gut microbiota profiling to examine social associations, space use, and microbiota similarity among individuals of three sympatric rodent species - wood mice (*Apodemus sylvaticus*), yellow-necked mice (*A. flavicollis*) and bank voles (*Myodes glareolus*), in two woodlands. By integrating multi-species social networks and spatial overlap with individual microbiota data, we test how both conspecific and interspecific interactions shape gut microbiota composition.

**Results:** At both study sites, social associations strongly predicted microbiota similarity among conspecific individuals, whereas spatial proximity consistently predicted microbiota similarity among interspecific pairs. In addition, social associations between mice of different species predicted microbiota similarity, although this effect was weaker than that observed within species and was asymmetric, with evidence for stronger microbial transmission from wood mice to yellow-necked mice than vice versa. By focusing on species-indicative bacterial taxa, we further show that wood mice with more social connections to yellow-necked mice harboured a greater richness of gut bacteria typical of yellow-necked mice.

**Conclusions:** Together, these results demonstrate that individual-level social behaviour can influence gut microbial transmission across species boundaries, particularly among closely related species with overlapping ecological niches. Interspecific social interactions may therefore represent an underappreciated pathway for the acquisition of novel gut symbionts in multi-species communities.

## Introduction

The gut microbiota is increasingly recognised as a key modulator of animal biology, with roles in digestion ^1–3^, immune regulation ^4,5^, energy homeostasis, ^6–8^, and behaviour ^9–11^. Yet gut microbial communities in many animals vary enormously among individuals and over time. For many animal species, transmission of microbes through conspecific social contacts is a key factor shaping microbiome variation. For example, sharing the same living environment has been shown to homogenise the gut microbiota of cohabiting humans ^12,13^, laboratory rodents ^14,15^, domestic animals ^16,17^ and wild animals such as baboons and horses ^18–20^. Pairwise social contacts have been shown to predict gut microbiota similarity in various social ^21–23^, semi-social ^24,25^ and even relatively asocial mammals ^26^, with some studies showing these effects are independent of genetic relatedness ^21,24^ or spatial overlap among interacting individuals ^24,25^. Similarly, social relationships among humans have been shown to predict gut microbial strain sharing ^27^, with more intimate relationships predicting more shared microbial strains ^28^. Importantly, such social influences or cohousing effects often outweigh the influence of individual-level factors in explaining microbiota composition within species ^18,21,22,24,29^.

Because of the key role of transmission in shaping host-associated microbial communities, microbiotas are usefully conceptualised as metacommunities - that is, as networks of local microbial communities inside individual hosts that are connected by microbial dispersal among host individuals through their contact (Miller et al., 2018; Miller & Bohannan, 2019; Robinson et al., 2019). Traditional metacommunity theory predicts that in patch-like ecological communities like islands 1) pairwise connectivity among communities should increase their similarity and that 2) local species diversity in a community should increase with its connectivity to others. These predictions often hold true in microbiome metacommunities: social association strength (pairwise connectivity) has been linked with more similar gut microbiotas among interacting humans ^27,28,33,34^, wild primates ^21–23^ and wild rodents ^24,25^. Similarly, the number of social connections an individual has (overall connectivity) has been shown to positively predict microbiota alpha diversity in humans ^35^, other primates ^23^ and rodents ^24^. However, almost all studies of social transmission of microbiota focus on sociality among members of the same species. Far less is known about the transmission processes occurring between individuals of different species. In a rare study exploring effects of cohabitation on the gut microbiota, Song et al. showed that humans share more skin microbes with their own dogs than other dogs ^13^. However, little is known about the effect of cross-species social behaviour on gut microbiota transmission and it remains unknown to what extent direct contacts among individuals of different animal species may function as routes of commensal microbiota transmission.

Although studies on social microbiota transmission focus on a single host species, these populations typically exist in larger mixed-species communities in which heterospecific individuals commonly interact. These heterospecific pairs of individuals, and especially those with strongly overlapping niches, may share gut microbes directly through social contacts (e.g., aggressive encounters or affiliative interactions like grooming, food sharing) or indirectly through shared environments (e.g., faecal contact at shared feeding or nesting sites). The social interactions occurring among heterospecific individuals likely differ from those between conspecifics, which could generate qualitatively and quantitatively different signatures of social transmission. For example, social behaviours involved in pair-bonding or reproduction (e.g., grooming, licking) will be rare among members of different species, while the contribution of indirect contact behaviours (e.g., coprophagy or encounters at shared feeding sites) to transmission may be greater among hetero-than conspecifics. Trophic relationships can also create transmission routes for the gut microbiota among species. For instance, while the gut microbiota of mammalian carnivores is compositionally distinct to that of herbivores ^36^, predatory mammals were shown to host more similar gut microbes to their common prey animals than to other sympatric herbivores ^37^.

Although mammalian species harbour species-characteristic gut microbiotas, these can converge to an extent in sympatry. For example, gut microbiota similarity among heterospecific individuals was shown to be higher among sympatric vs allopatric individuals, in small mammals and primates ^18,38–40^. Similarly, gorillas and chimpanzees inhabiting the same geographic area shared more gut microbes than gorillas and chimpanzees living far apart ^39^. Further, in a study comparing gut microbiota of multiple mammalian species across the whole western hemisphere, sympatric populations of different species shared more microbiota than allopatric ^37^.

Such convergence in the microbiota of different species in sympatry may be a result of interspecific social transmission, but other processes could also be involved including trophic interactions, shared exposure to the same environmental pools of microbes, or similar selection pressures on the microbiota when living in the same place, for example through shared diet. However, to date no studies of wild animals have resolved patterns of cross-species microbiota transmission at an individual-level. Neither do we know to what extent microbiota similarity between sympatric individuals arises through direct (e.g., through physical contact) vs. indirect (e.g., through the shared environment) microbial transmission. Teasing apart these effects is challenging, as it requires fine-scale individual level behavioural data from many individuals of multiple sympatric species. Specifically, to answer questions about how different forms of interaction may influence heterospecific microbiota transmission, and whether transmission processes are symmetrical or asymmetrical (driven more by transmission from one species to the other), detailed high coverage data on social networks and spatial distributions of individuals of multiple coexisting species are needed.

Here, we address this knowledge gap by using RFID technology to intensively track social interactions and space use among individuals of three sympatric wild rodent species at two distinct woodland sites, while collecting parallel data on their gut microbiota. At both study sites, previous work has indicated strong effects of conspecific social microbiota transmission among wood mice (*Apodemus sylvaticus*) ^24,25^. Here, we extend this to consider multispecies rodent communities at both sites, including yellow-necked mice (*Apodemus flavicollis*) and bank voles (*Myodes glaerolus*) as well as wood mice. Using this data, we assess how social and spatial associations among rodents predict gut microbial sharing among both con- and heterospecific pairs of individuals. Specifically, we test three predictions about cross-species sharing of gut microbes:

1. First, we predict that both direct (via social contact) and indirect (via shared environments) mechanisms could drive gut microbial transmission among heterospecifics. However, as animals usually interact more intimately with individuals of their own species, evidence for transmission via social contact should be weaker among heterospecifics than conspecifics. Transmission by social contact is expected to generate a positive correlation between heterospecific social association and gut microbiota similarity, that is distinct from any effect of spatial proximity alone. Conversely, indirect transmission is expected to generate a positive relationship between spatial proximity to heterospecifics and gut microbiota similarity, that is distinct from any effect of social association. Overall, therefore, we predict positive and distinct effects of social association with, and spatial proximity to heterospecifics, on pairwise microbiota similarity among heterospecifics.
2. Second, we predict that (under the expectation of phylosymbiosis and each host species harbouring some species-indicative microbial taxa that are enriched in that species), individuals most strongly socially connected to members of another species should harbour a greater diversity of that other species’ indicative microbial taxa.
3. Third, because a degree of species distinctness in the microbiota is expected, we predict a positive relationship between an individual’s overall social connectedness to heterospecifics and their microbiota richness.

By testing these three predictions, we aim to create more nuanced understanding of the role of social behaviour between members of different species in spreading gut microbes across species boundaries.

## 5.2 Methods

### Study sites and field data collection

At two independent study sites, we studied communities of three common woodland rodent species that frequently co-occur across Europe: wood mice (*Apodemus sylvaticus*, AS), yellow-necked mice (*Apodemus flavicollis,* AF) and bank voles (*Myodes glareolus,* MG). The study sites were 40 miles apart, at Holly Hill, Wytham Woods in Oxfordshire (“Wytham”, 160 x 160 m, 51.77 lat, −1.33 long) and Nash’s Copse, Silwood Park in Berkshire (“Silwood”, 200 x 120 m 51.41 lat, −0.64 long), in the UK. At each site these species have overlapping home ranges and partially overlapping diets, and nest in underground burrows. They engage in a range of ecological and social interactions that could facilitate gut microbe transmission, including i) shared exposure to environmental microbes (e.g. from soil, diet or vegetation) wherever home ranges overlap, ii) exposure to gut microbes through social contact behaviours such as aggression ^41–43^ and iii) indirect exposure to gut microbes through exposure to faeces in the environment.

On a ∼2.5ha plot at each study site, we used a combination of regular trapping, PIT-tagging and PIT-tag loggers to collect high-resolution spatiotemporal data on individual whereabouts. Species were present in similar relative abundances at both sites, with wood mice dominating in numbers. At both sites trapping was carried out approximately fortnightly, over a 10-month period in Wytham and a 12-month period in Silwood). At first capture, all rodents were injected subcutaneously with a passive integrated transponder (PIT)-tag for permanent identification and tracking. At each trapping, demographic characteristics (age and sex) were recorded, and faecal samples for gut microbiota analysis (∼50-300 mg) collected from traps with sterilized tweezers and frozen at −80°C within 12 hours of collection. Traps were sterilized by washing in bleach solution between trappings (with additional autoclave of traps in Silwood) to avoid cross-contamination of samples. Rodents were tracked throughout each period using custom-build PIT-tag loggers - devices that record the time-stamped presence of any tagged rodent within the detection range of the coil. Loggers were evenly spaced across each monitoring grid and rotated around the study area to enable even spatial coverage in rodent monitoring. While the general principles of data collection were comparable between sites, the study areas, logger design and logger rotation protocols differed somewhat between the two sites, with an improved logger design and greatly increased logger density in Wytham (n=60 loggers in Wytham, n=9 loggers in Silwood, each across a ∼2.5ha plot; see comparisons of the two data sets in Table S1, Figure S1). However, both methodologies resulted in even spatial coverage of the monitoring area over the study period, allowing robust estimation of social networks and space use.

### Describing social and spatial behaviour of rodents

We used spatiotemporal co-occurrence data from RFID loggers to build individual-by-individual association matrices describing both social networks and spatial proximity among all sympatric rodent individuals, at each of the two sites. Social networks and spatial proximity matrices were built using the same methods as in ^25^, as described briefly below. Only unlike in our previous work focusing solely on wood mice, here association matrices were constructed using data from all tagged rodents that were recorded by loggers in each site. All analyses were conducted in R ^44^.

### Social network construction

Following the principles of animal social network analysis ^45,46^. individuals were classified as nodes in the social network, with edges representing associations between individuals and edge weight based on the frequency of instances when two individuals were observed ‘associated’ relative to the frequency they were observed overall. Here, ‘associated’ refers to two individuals that were observed within 12 hours of each other at the same logger location during the same night, defined as the 16 hour period from 4pm to 6am, the daily period when all three rodent species are active (Figure S2). From the raw counts of co-occurrence incidents, social association matrices were derived by calculating the life-span overlap corrected simple ratio index (“Adjusted SRI”) ^24,47^. Individual ‘lifespans’ were taken as the time between their first and last logger observation. This association index was defined as:

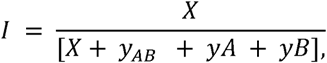

where *X* is the number of nights in which individuals *A* and *B* were observed associated (at the same logger location within 12 h of each other), *y_AB_* is the number of nights in which *A* and *B* were both observed but not associated (observed at different logger locations, or at the same location but more than 12□h apart) and *y_A_* and *y_B_* are the number of nights in which both were known to be alive but only *A* or *B* was observed, respectively. Accounting for lifespan overlap in this metric allows us to more accurately summarize the temporally fluctuating social structure of the rodent population in one static social network. While this type of measure of spatiotemporal co-occurrence is not a measure of direct social interaction, it is a commonly used and powerful proxy for actual social relationships ^24,25,47–51^.

### Home range overlap and spatial proximity estimation

To derive a matrix reflecting pairwise degrees of spatial overlap among individuals, we first estimated home ranges as 75% core of the kernel density utilization distribution of each individual’s tracking records using the R package ctmm, and used the Bhattacaryya coefficient to estimate overlap between any two ranges (See Supplementary Appendix S1 for details on this method). However, as this method of estimating spatial overlap is data hungry, accurate home ranges could only be estimated for individuals with sufficient logger records (only Wytham rodents, 52% of AF, 81% of AS, 73% of MG), that is with more than five unique observations in more than three unique logger locations which were statistically indistinguishable from random draws from a stable range (i.e., satisfied the “variogram criteria” described in Supplementary Appendix S1). We therefore adopted a simpler metric of *spatial proximity* that could be calculated for all pairs of individuals, based on the distance between the centroids of their location records. Metric distances between centroids were scaled between 0 and 1 and the complement taken to derive a measure of spatial proximity (spatial proximity = 1-scaled distance). Among pairs of individuals where both measures could be derived, spatial proximity measure correlated well with home range overlap (Bhattacaryya coefficients) up to ∼120m of centroid distance (Pearson’s correlation coefficient= 0.66, p<0.01, Supplementary Appendix S1, Figure S3). Thus, we used spatial proximity as a proxy for home range overlap for animals whose centroids were up to 120 meters apart, and set spatial overlap to 0 for those with centroid distance >120m apart. This reflects the range sizes detected in our data, with larger ranges reaching areas over 8000 m^2^. Thus, generalising the ranges as circular, the centroid distance where no ranges should overlap should be over 100 meters.

### Estimating an individual’s connectedness to others through social contacts and shared space

In order to test our second and third predictions on how an individual’s relative exposure to heterospecific individuals influences their microbiota, we calculated node-level metrics reflecting each individual’s level of social connectedness to conspecific, and heterospecific individuals in the social network (i.e. measures of social centrality). For these analyses we used wood mice as a focal species as they were the most abundant species and the only species with sufficient sample size for robust analyses of this type at both sites (Table S1C). First, we calculated each wood mouse individual’s conspecific and heterospecific degree (CD and HD), that is the number of social connections they had to individuals of their own species, or other species, respectively. Second, we calculated species-specific heterospecific degrees: AF-degree (number of connections to *A. flavicollis*) and MG-degree (number of connections to *M. glareolus*). And finally, we calculated a set of custom metrics that capture not only first-order connections to heterospecific individuals, but also second-order connections through other wood mice. This measure which we term *extended heterospecific degree* (Extended HD) was defined as:

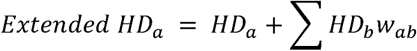

where *HD*_a_ is individual a’s heterospecific degree and ∑*HD*_a_*W*_ab_ is the sum of the heterospecific degrees of all direct contacts of individual a (*HD*_b_) multiplied by the social association strength between individual a and each direct contact b respectively (*W*_ab_). Extended heterospecific degree was also calculated for all wood mice on a species-specific basis, to derive measures of each individual’s connectedness to yellow-necked mice (“Extended AF-degree”) and bank voles (“Extended MG-degree”).

Finally, in addition to these measures of social centrality, we also derived a parallel set of node-level ‘spatial density’ metrics to capture each individual wood mouse’s range overlap with other rodents. For this, we used the spatial proximity matrix to calculate the spatial density of each type of other rodent around each wood mouse’s location centroid. Spatial density was thus calculated by summing each wood mouse’s spatial proximity values to their conspecifics or heterospecific contacts respectively. This yielded individual-level density measures for conspecifics (“Conspecific density”), heterospecifics (“Heterospecific density”), yellow-necked mice (“AF density”) and bank voles (“MG density”).

### Laboratory work and bioinformatics

We successfully characterised the gut microbiota from 405 faecal samples belonging to 217 rodent individuals in Wytham (AS: 362 samples from 189 individuals, AF: 21 samples from 15 individuals, MG: 22 samples from 13 individuals) and from 269 samples belonging to 88 individuals in Silwood (AS: 253 samples from 75 individuals, AF: 32 samples from 13 individuals). No bank vole microbiota samples from Silwood were collected, and thus all our data on vole microbiota comes from Wytham. The gut microbiota was profiled with amplicon sequencing of the 16S rRNA marker gene (V4 or V4-5 region), sequenced on the Illumina MiSeq platform. Samples from the two sites were processed and analysed similarly, except a different reverse primer that yielded slightly longer 16S sequences was used in Wytham. Full details of library preparation, sequencing and bioinformatics are given in Supplementary Appendix S2 with methodological differences between the two data sets summarised in Table S1. For both data sets, sequence data were processed through the DADA2 pipeline (Callahan et al., 2016; versions 1.6 and 1.14), to infer amplicon sequence variants (ASVs). Using package *phyloseq* ^53^, ASV counts were normalized to proportional abundance within each sample ^54^ and singleton ASVs as well as those belonging to non-gut microbial taxa (Cyanobacteria, Mitochondria, Xanthomonadales) were removed. Lastly, we used sample completeness and rarefaction curves made with package *iNEXT* ^55^ to decide read depth thresholds below which samples were dropped from the dataset (6,000 in Wytham and 10,000 in Silwood). Despite no visible bands in negative controls when run on a gel, the Wytham dataset had detectable levels of contamination in sequenced negative controls, which we dealt with by removing taxa labelled as contaminants using the *decontam* algorithm ^56^, and by including extraction/PCR plate (the two are equivalent in this dataset) as a technical covariate in all models. For consistency, extraction/PCR plate was also included as a covariate in models using Silwood data.

### Describing the gut microbiota of woodland rodents

To characterise variation in microbiota diversity and composition among rodents, we used presence-absence –based, rather than abundance-weighted alpha and beta diversity metrics as previous work in this study system showed microbiota transmission signals were more detectable using presence/absence variation rather than abundance variation ^25^, likely because transmission primarily influences which taxa end up in hosts, while taxon abundances may be more influenced by selective forces shaping microbial growth within hosts. Thus, to estimate microbiota alpha diversity, we estimated microbiota richness (Observed richness in a sample, after filtering out singletons, doubletons and non-gut microbial taxa), and to characterise beta diversity (similarity), we calculated the Jaccard Index (the proportion of shared ASVs) for each sample pair. Read depth was used as a control variable in all models predicting alpha diversity to account for any sampling bias in observed richness. To estimate the extent to which individual wood mice harboured microbial taxa (ASVs) characteristic of another host species (AF or MG), we first utilised Random Forest Classifier (RFC) algorithms ^57^ implemented in package *randomForest* ^58^, to identify ASVs that were indicative of each host species, i.e., those with high importance in allowing the algorithm to predict host species from microbiota data. Each RFC model included wood mice and one other species (AF or MG) and was performed with 1500 trees and repeated 3 times, pruning away taxa with importance values of zero between each run. Taxon importance in each RFC model was assessed using the mean Decrease in Accuracy (the mean change in out-of-bag error estimates when data for that ASV was permuted). We considered AF or MG-indicator taxa to be those with i) importance values >0 in the final RFC model and ii) higher relative abundance in that species compared to wood mice. Thus AF-indicator taxa were ASVs that were important for distinguishing AF from AS and enriched in AF compared to AS, while MG-indicator taxa were ASVs that were important for distinguishing MG from AS and enriched in MG compared to AS. For each wood mouse, we then calculated the number of species-indicator taxa (AF-indicator or MG-indicator richness). Finally, to serve as a null comparison to trends related to AF-indicator richness, we also derived a complementary set of AS-indicator taxa (those important in distinguishing AS from AF and enriched in AS compared to AF) and calculated the richness of these “own-species” indicators in each wood mouse.

### Statistical analyses

#### Estimating host species specificity in microbiota composition

To test for species differences in microbiota composition we used a marginal PERMANOVA with Jaccard Index as the response, implemented with the adonis2 function in package vegan ^59^. Models were run separately for Wytham and Silwood and host species specificity of the microbiota was visualized using Principal Coordinates Analysis (PCoA).

#### Social and spatial effects on microbiota similarity

To test whether microbiota similarity among conspecific vs. heterospecific pairs rodents was predicted by spatial proximity or social association strength, we performed Bayesian multi-membership beta-regression in package *brms* ^60^, as described previously (Raulo et al., 2021 and Raulo et al. 2024). Here, we focused only on the conspecific and heterospecific associations of our focal species the wood mouse (the most numerous species in the data), and thus our model only considered sample pairs involving wood mice (AS-AS, AS-AF and AS-MG pairs). Models used Jaccard Index of microbiota similarity as the response variable, excluding comparisons of samples from the same individual. As main predictors of interest, we fitted social association (Adjusted SRI) and its interaction with the species combination of a pair (3-way factor: AS-AS, AS-AF or AS-MG). Similarly, we fitted spatial proximity and its interaction with species combination. To control for potential confounding variables that could drive an effect of social or spatial proximity on microbiota similarity, we fitted several dyadic control variables: similarity in (age and sex) (same or different), similarity in age class (same or different) and sampling interval (time in days between which samples were taken, scaled between 0 and 1). We also fitted technical covariates: difference in sample read depth (scaled between 0 and 1), PCR plate similarity (same or different), and for Wytham data also extraction plate similarity (same or different). To control for non-independence among pairwise measures, the intercept was allowed to vary as defined by two multi-membership random effects, one capturing the individuals (Individual A + Individual B), and another capturing the samples (Sample A + Sample B) involved in each sample pair. This random effect structure controls both for the inherent non-independence of the dyadic response variable and the autocorrelation arising from the presence of repeat microbiota measurements for some individuals in the dataset.

#### Posthoc models for asymmetric social effects on microbiota similarity

The above described pairwise model used symmetrical data, with each pair of samples represented by one measure of microbiota similarity, social association, spatial proximity etc. To explore the directionality in identified transmission effects we also used an asymmetric version of this model. These models were used to test whether microbiota homogenisation associated with heterospecific social associations is generated by equal bidirectional transmission between species or whether the microbiota of one host species is more invasive to the other than vice versa. For this we calculated *asymmetric Jaccard microbiota similarity* measures as well as *asymmetric social association indices* for all AS-AF sample pairs. Asymmetric Jaccard similarities were calculated as the number of microbial taxa shared by samples A and B divided by the total number of taxa in sample A (Jaccard similarity in one direction) or B (Jaccard similarity in the other direction). Similarly, the asymmetric social association index was calculated as in the general SRI described above, but with each pair getting two measures of social association: the instances A and B were observed associated divided by the total number of instances individual A was observed with or without B (social association in one direction) or the total number of instances B was observed with or without A (social association in the other direction). These asymmetric measures are akin to the common logic of directional edges in social network analysis ^61^ although here instead of commonly used measures like counts of instances (e.g., how many times individual A groomed individual B) we use a proportional index (how many times individual A was observed with individual B, out of all the times individual A was observed). Our asymmetric measures of social association and microbiota similarity were then included in a Bayesian multi-membership model (similar to above) to explore whether an observed effect of AS-AF social association on microbiota similarity was more driven by transmission from AS to AF or AF to AS individuals. Specifically, we ran two models using the same set of AS-AF sample pairs, but with the first model including the variables in one direction (AS to AF) and the second model in the other direction (AF to AS).

#### Social and spatial effects on indicator taxon richness

To test our second prediction (that heterospecific social centrality or spatial density will predict the richness of heterospecific species-indicator gut microbes) we used Bayesian Poisson regression models, implemented in R package *MCMCglmm* ^62^. We ran two models for wood mice, one predicting AF indicator richness with AF social degree and AF density, and another predicting MG indicator richness with MG social degree and MG density. In each model, conspecific social degree, conspecific density, month, sex, age class, read depth and total microbiota richness were included as covariates, as well as random effects for PCR plate and individual ID. Including total microbiota richness as a covariate ensures that any effect of heterospecific social centrality or spatial density is not merely an effect on overall microbiota richness, but relates specifically to the richness of species-indicator taxa. As a control comparison (for which we would not expect to see an effect of heterospecific social centrality or spatial density), we also ran a similar model predicting the richness of own-species indicator taxa (AS-indicator richness) as a function of both their social and spatial connectedness to both heterospecific (AF) and conspecific individuals.

#### Social and spatial effects on overall microbiota alpha diversity

To test our third prediction, we used a Poisson regression model similar to above but with overall microbiota richness (the observed number of ASVs after filtering, see above) as the response, conspecific and heterospecific degree and spatial density as predictors, as well as the same covariates and random effects as for the species indicator richness models described above.

## 5.3 Results

### Social structure and spatiotemporal distribution of sympatric rodent species

We followed the behaviour of 203 rodents in Wytham (150 AS, 18 AF, 35 MG) and 159 in Silwood (83 AS, 26 AF, 49 MG) over a 10- and 12-month period respectively. Multi-species social networks at each site (Figure 1 A-B) showed low but statistically significant social assortment by species i.e., individuals associated slightly more strongly with conspecifics than heterospecifics (network assortativity coefficient 0.10 in Wytham, 0.01 in Silwood). Social connectedness levels were generally comparable across the three study species, with slight species differences: yellow-necked mice tended to have more social connections (higher total degree), and bank voles fewer social connections than wood mice (Figure S4A). Home ranges were larger at Silwood than Wytham, and varied by species within each site, with mice having larger average home range size than voles, and yellow-necked mice having slightly larger home ranges than wood mice (Figure S4B). All three rodent species shared living space with each other (home ranges of heterospecific individuals frequently overlapped) and individuals of all sympatric species were frequently detected at the same logger within the same hour (Figure S2). The timing of these co-occurrences followed the general activity patterns of each species, with heterospecific co-occurrences generally happening during the time of the day when all three rodent species were active. For our focal species, the wood mouse, measures of heterospecific and conspecific degree were significantly positively correlated implying that wood mice who had social contacts with many other wood mice also tended to have more contacts with yellow-necked mice and bank voles (Figure S5).

**Figure 1.**
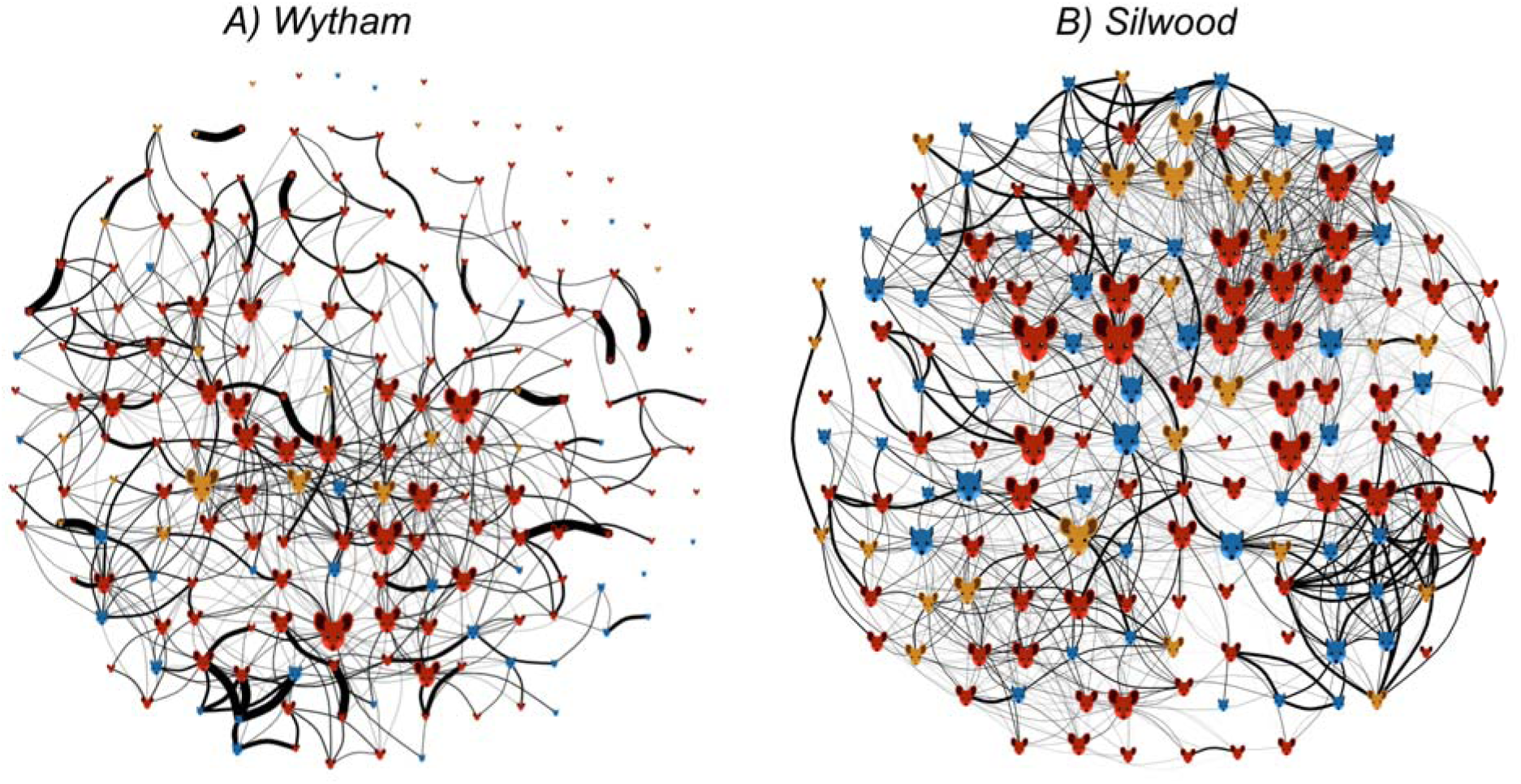
Multi-species social network of rodents in A) Wytham and B) Silwood. Nodes are individual wood mice (red), yellow-necked mice (yellow) and bank voles (blue) and edges are measures of social association (Adjusted SRI, based on the number of co-occurrences at an RFID logger on the same night). Node size depicts the overall degree of an individual, i.e., the number of social connections (larger = more connections). Line thickness denotes social association between individuals (thicker = higher SRI value).

### Microbiota differences among host species

At both study sites, microbiota composition differed significantly between the two *Apodemus* mouse species (PERMANOVA on Jaccard distance, AS vs AF: Wytham R^2^=0.02, F=2.05 p<0.01, Silwood R^2^=0.02, F=1.73, p=0.01), though clustering of the two mouse species was not obvious on the first two axes in a principal coordinates analysis (Figure 2). In Wytham, where data on vole microbiota was available, differences in mouse and vole microbiota composition were greater than those among the two species of mice (PERMANOVA on Jaccard distance, AS vs MG in Wytham: R^2^=0.05, F=10.53, p<0.01).

**Figure 2.**
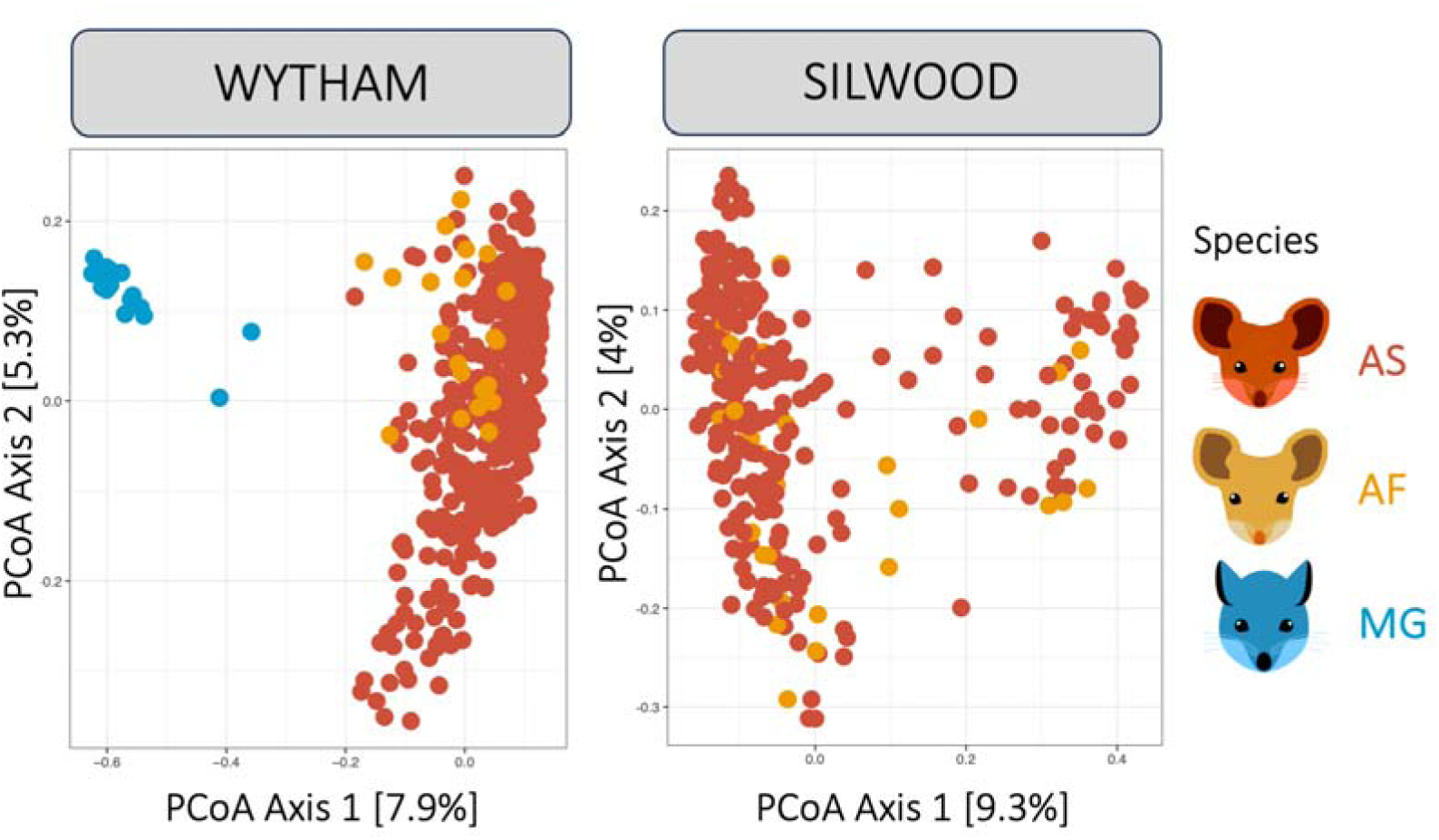
PCoA clustering of gut microbiota composition according to host species (colour) at the two study sites, Wytham and Silwood.

### Social and spatial effects on microbiota similarity

In both study sites, effects of social association and spatial proximity on microbiota similarity varied significantly between conspecific and heterospecific pairs (interactions between social or spatial terms and species combination of the pair; Tables S2A-B). Overall, gut microbiota similarity (Jaccard Index, the proportion of shared taxa) was predicted by both social association with and spatial proximity to conspecifics, while only spatial proximity to heterospecific individuals (and not heterospecific social association) consistently predicted microbiota similarity (See Tables S2A-B, Figure 3). Specifically, among conspecific wood mouse pairs (AS-AS), social association strongly and positively predicted microbiota similarity (Beta-regression, Wytham: Posterior mean 0.26, CI=0.21 to 0.31; Silwood: Posterior mean 0.52, CI=0.48 to 0.56), while spatial proximity had a much weaker yet significant effect on microbiota similarity (Betaregression, Wytham: Posterior mean 0.06, CI=0.05 to 0.06; Silwood: Posterior mean 0.03, CI=0.02 to 0.04; Figure 3). Conversely, among heterospecific pairs of mice (AS-AF), microbiota similarity was predicted by individuals’ spatial proximity at both sites (Betaregression, Wytham: Posterior mean 0.05, CI=0.03 to 0.06; Silwood: Posterior mean 0.04, CI=0.02 to 0.06), and also by their social association at one site (Silwood; Betaregression Posterior mean 0.27, CI=0.20 to 0.35), though this effect was smaller than that of conspecific social association (Figure 3). Between wood mice and bank voles (AS-MG pairs), neither spatial nor social proximity significantly predicted microbiota similarity (Figure 3).

**Figure 3.**
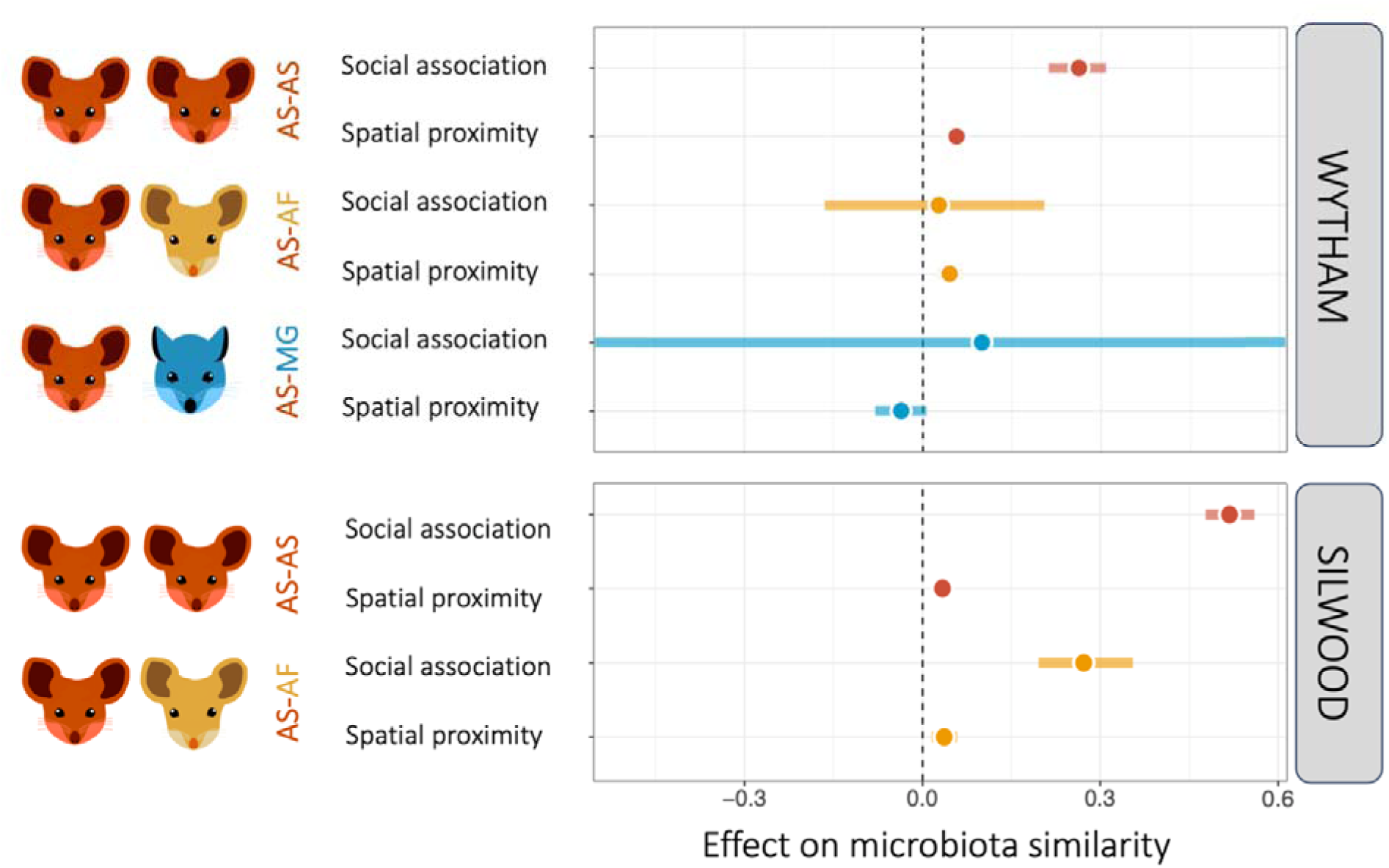
Effects of social association and spatial proximity on microbiota similarity between conspecific (AS-AS) and heterospecific (AS-AF & AS-MG) pairs of sympatric rodents at the two study sites, Wytham and Silwood. Effect size estimates (points) and their 95% credible intervals are plotted from Bayesian multi-membership beta regression models with pairwise microbiota similarity among hosts (Jaccard Index) as the response and an interaction term between social association*species or spatial proximity*species as main predictors (model results in Tables S2A-B). Where credible intervals do not overlap zero, a variable significantly predicts microbiota similarity. The credible intervals of AS-MG social association effect here (Posterior mean −0,26, CI= −1.9 to 1.3) extend beyond the x axis boundaries.

We explored the observed effect of heterospecific social association on microbiota sharing between AS-AF pairs in Silwood further with a set of posthoc models disentangling the effects of AS-to-AF and AF-to-AS social transmission. These revealed asymmetric effects of AS-AF social association on the microbiota of each species, with a stronger influence of wood mice on the microbiota of yellow-necked mice than vice versa (See Tables S3A-B, Figure 4). In other words, when a yellow-necked mouse was socially associated with a given wood mouse, the proportion of its microbiota shared with that wood mouse was larger than the proportion the wood mouse shared with the yellow-necked mouse (Betaregression, effect of AS-to-AF social association on asymmetrical microbiota similarity: Posterior mean 0.10, CI=0.05 to 0.16; Table S3A & effect of AF-to-AS social association on asymmetrical microbiota similarity: Posterior mean 0.27, CI=0.21 to 0.33; Table S3B).

**Figure 4.**
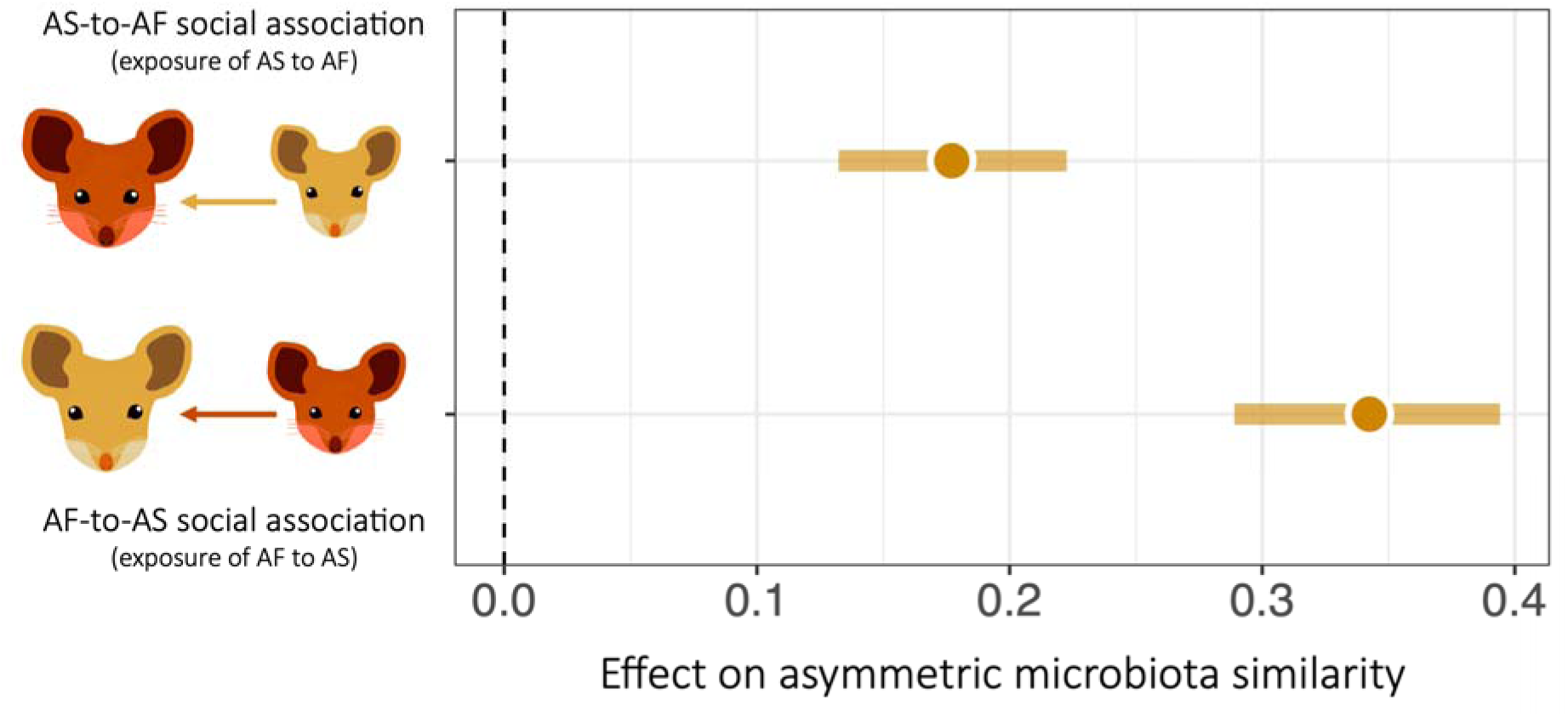
Effects of asymmetric social association on asymmetric microbiota similarity among AS-AF pairs in Silwood. Effect size estimates (points) and their 95% credible intervals (coloured lines) are plotted from the Bayesian multi-membership beta regression models with asymmetric microbiota similarity (asymmetric Jaccard Index) as the response and asymmetric social association as the main predictor (Model results in Tables S3A-B). For AS-to-AF social association, asymmetric microbiota similarity is the proportion of taxa in the AF individual that were also observed in the AS individual; for AF-to-AS social association, asymmetric microbiota similarity is the proportion of taxa in the AS individual that were also observed in the AF individual. Where credible intervals do not overlap zero, a variable significantly predicts microbiota similarity and where credible intervals do not overlap each other the effects are significantly different.

### Social and spatial effects on indicator taxa

At both study sites, the social connectedness of wood mice to yellow-necked mice (AF-degree) positively predicted the richness of AF-indicator taxa in their gut (Poisson regression, Wytham: Posterior mean 0.19, CI=0.07 to 0.33; Silwood: Posterior mean 0.19, CI=0.14 to 0.36); Table S4A-B, Figure 5). These effects were statistically significant even when controlling for social connectedness to other wood mice (conspecific degree), spatial density of yellow-necked mice, spatial density of conspecifics and other covariates. By contrast, AF-degree did not positively predict the richness of AS indicator taxa and in fact this relationship was negative at both sites (Figure 5; Poisson regression, Wytham: Posterior mean −0.07, CI=-0.12 to −0.01; Silwood: Posterior mean −0.12, CI=-0.18 to −0.05); Tables S6A-B). AS indicator richness tended to be positively predicted by conspecific degree, though this effect was only significant in Silwood (Poisson regression, Wytham: Posterior mean 0.05, CI=-0.01 to 0.12; Silwood: Posterior mean 0.14, CI=0.09 to 0.22); Tables S6A-B; Figure 5). The positive effects of AF-degree on AF indicator richness remained of a similar magnitude whether using AF-degree or extended AF-degree as a predictor at both sites (Table S5A-B), suggesting second-order connectedness of wood mice to yellow-necked mice had no additional effect on the wood mouse microbiota. Finally, while no measure of social connectedness to bank voles significantly predicted MG indicator richness in wood mice, MG indicators were significantly enriched by spatial density of bank voles around the location centroids of wood mice in Wytham (Poisson regression for Wytham: posterior mean 0.47, CI=0.07 to 1.28; Table S7A). Spatial density of yellow-necked mice had no effect on AF indicator richness in either population.

**Figure 5.**
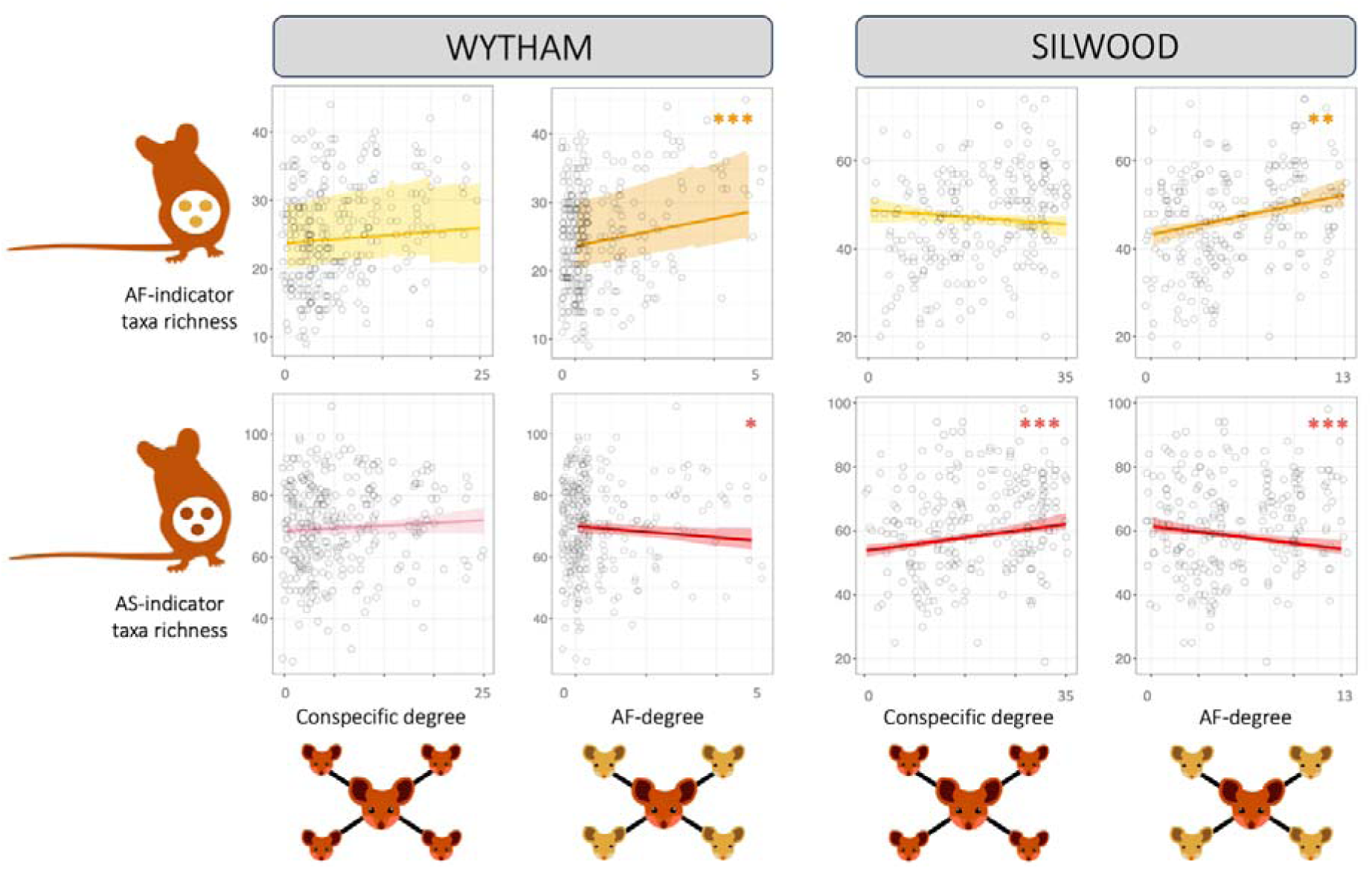
Effects of conspecific and heterospecific social connectedness on host species indicative taxa in the gut microbiota of wood mice. Estimated slopes and their 95% credible intervals from Bayesian Poisson regression models predicting (AF or AS) indicator taxa richness with conspecific or AF-degree. Slopes are depicted together with raw data points (circles) (Model results in Tables S4 & S6). Significant trends are marked with stars and darker coloured lines.

### Social and spatial effects on overall microbiota alpha diversity

No measure of social connectedness to other species predicted overall microbiota richness for wood mice at either study site. However, in Silwood, heterospecific density around the centroid of a wood mouse’s spatial location positively predicted overall microbiota richness (Poisson regression, Silwood: Posterior mean 0.21, CI=0.03 to 0.38; Table S8B), while conspecific density had a negative effect on microbiota richness (Poisson regression, Silwood: Posterior mean −0.30, CI=-0.47 to −0.12; Table S8B). No effects of spatial density were detected in Wytham.

## 5.4 Discussion

Through individual tracking in mixed-species wild rodent communities at two study sites, we present the first evidence that individual-level interactions among members of different species shape cross-species gut microbial transmission among sympatric wild mammals. While other research has emphasized that sympatric populations of different host species may harbour more similar microbiota than allopatric populations ^37–40^, whether this pattern arises through convergent exposure to the same environments or more direct exposure to heterospecific individuals has so far not been directly investigated. We provide evidence that individual-level variation in social behaviour and space use can drive gut microbial transmission across species boundaries, and that individuals with stronger transmission links to heterospecifics tend to harbour more of those species’ characteristic gut microbes. Shared use of space consistently homogenized the gut microbiota of heterospecific individuals as much as it did for conspecifics. Meanwhile social association had a strong homogenizing effect on overall microbiota composition among conspecifics, but a weaker or undetectable effect among heterospecifics. Despite this, our results did indicate that for wood mice, being socially connected to many yellow-necked mice did in fact introduce yellow-necked mouse – typical bacteria into their microbiota. This effect was consistent at both our study sites and was not dependent simply on the extent of space sharing with yellow-necked mice, suggesting a more direct transmission link than exposure to the same environmental pool of microbes. These results emphasize the importance of considering high-resolution animal contact networks, both within and among species, for understanding the impact of transmission on microbiota composition.

Interspecific transmission of gut microbes between the two mouse species revealed here could occur through a range of contact behaviours, from direct aggressive encounters to indirect exposure to faeces while sharing burrows. Encounter experiments, whereby *<u>Apodemus</u>* mice of different species are temporarily introduced to one another in a cage, have implied that yellow-necked mice (the larger species) are socially dominant over wood mice ^41,42^ and that individuals of both species behave more agonistically towards each other than towards conspecifics ^41^. However, severe aggression was still rare and more amicable social behaviours, such as nose-to-nose contact and allogrooming have also been observed between wood mice and yellow-necked mice ^43^. Higher connectedness to heterospecifics might also affect stress levels, which could in turn affect the microbiota ^63–65^. However, hormonal changes could not explain the finding that connectedness to a specific species predicted enrichment of the gut microbiota by specific indicator taxa of that species, which constitutes strong evidence for direct social transmission of gut microbes across species boundaries.

Interestingly, in the study populations where we found social association between wood mice and yellow-necked mice did homogenise their overall gut microbiota profiles, this effect was not symmetrical on each species. Yellow-necked mice converged on the gut microbiota composition of their wood mouse contacts more strongly than wood mice converged on the microbiota of their yellow-necked mouse contacts. To our knowledge this is the first report of asymmetric microbiota transmission among wild animals. Somewhat similar patterns were observed in humans and their pet dogs in a study that found introducing a dog to a family resulted in the human gut microbiota converging on the dog microbiota much more than the dog’s microbiota was influenced by the humans’ microbiota ^13^. The asymmetric transmission effects we observe may arise from asymmetric behavioural interactions between species (e.g. biting), differences in transmissibility or invasiveness among microbial taxa, or variation in how susceptible each host species microbiota is to invasion. To disentangle these influences, useful next steps include mixed-species co-housing experiments, in which microbiota-transmitting behaviours are monitored in detail, and controlled inoculation experiments in which microbiotas from different species microbiotas are inoculated into laboratory mice in varying orders to quantify differences in their invasiveness across alternate resident microbiotas.

While gut microbes seem to be transmitted through some kind of behavioural contact between wood mice and yellow-necked mice, this was not true between wood mice and bank voles, whose microbiota was not homogenised by social associations. This lack of social transmission between wood mice and bank voles was not driven by fewer social associations between these two species, as wood mice had just as many and as strong associations with voles as with yellow-necked mice. In fact, earlier research found that bank voles were more often observed at a feeding site with wood mice than with other bank voles (Lambin, 1988). However, the two mouse species have more overlapping niches, such that social association (spatiotemporal co-occurrence) may be more likely to reflect physical interactions among them. Alternatively, a closely related sister species will possess a more similar gut physiology and immune system, which could facilitate colonization and persistence of mouse-typical taxa in wood mice more than vole-typical taxa. In line with this, earlier research indicated that gut microbiota sharing between sympatric species seems to be particularly influential between closely related species. For example, while many primate species are known to harbour a highly species-specific microbiota even when inhabiting the same environment ^67^, among populations of closely related baboon sub-species, spatial overlap was a stronger predictor of microbiota composition than subspecies identity ^18^.

Our results imply that social interactions with other heterospecific individuals can shift one’s microbiome to more closely resemble that of the other species. Further, we found that sharing space with heterospecifics tended to increase the gut microbial alpha diversity of our focal species. Such gut microbial exchanges between species through social interactions or space sharing hold potential for shaping larger scale biological processes. For instance, cross-species transmission of symbiotic microbes may result in horizontal transfer, or “lending”, of metabolic capacities, immune resilience, thermal tolerance or other microbiota-mediated traits between sympatric host species. This is analogous to animal hosts picking up useful microbial symbionts from their environment or diet, for example as bean bugs have gained insecticide resistance by taking up resistant soil microbes into their microbiome ^68^. Some evidence hinting towards the possibility of microbial transmission across species boundaries transferring new metabolic capabilities comes from research on the toxin-degrading microbiota of wood rats. Specific gut microbes present in some woodrat populations were shown to enable them to digest a tannin-rich plant ^1^ and experimental transplantation of this gut microbiota into domestic laboratory rats (a different species) enabled the latter to digest a tannin-rich diet that otherwise damaged their liver ^69^. The adaptive value of microbes transmitted naturally through social contacts between species might be especially high, as intimate contacts can spread strictly anaerobic symbionts that may be well adapted to a host-associated mutualist life style ^25,70^. Overall, microbiota sharing among sympatric species might shape species’ niches, making their social and ecological interactions potentially important drivers of their ability to adapt in changing or novel environments. For example, gut microbiota transmission among sympatric species could broaden their dietary niche, helping them to adapt to climate change induced dietary shifts. At the same time, picking up microbes from other species may also aid invasive species in adapting to novel environments ^71,72^. Indeed, to what extent invasive species’ acquisition of symbionts from local niche neighbours predicts their ability to invade niche space from local species is an important direction for future research. Acquisition of new metabolic capabilities through cross-species microbiota transmission might also shape species niches over longer evolutionary timescales. For example, many domesticated farm animals have evolved not only under human selection but also under continuous exposure to each other, and microbiota transfer between domestic species such as cows and dogs may have enhanced the latter’s ability to digest more plant-based diets while adapting from the ancestral carnivore diets of wolves to more omnivore diets of dogs living off the leftovers of humans ^73,74^. The effect of cross-species microbiota sharing on adaptive plasticity could be tested in future by comparing the microbiota and metabolic capabilities among populations of a given species living with or without closely interacting sympatric species. Identifying the types of microbes that are most prone to spread across species boundaries constitutes another key question for future research, including whether these are more or less likely to provide important functions, or conversely to have pathogenic potential.

Overall, this study provides clear evidence that individual-level social behaviours can influence the spread of gut microbiota between sympatric species, and highlights the value of combining individual-level tracking with a social network perspective to examine microbiota transmission dynamics. Understanding how microbiota spread across multiple levels of host organisation, from vertical transmission along maternal lines to horizontal transmission through social networks or ecological interaction networks of sympatric hosts, will help us sketch the shape of the ecological landscape in which these microbes live, move and evolve. Ultimately, this will enhance our understanding of the role host-associated microbiotas play in either linking or separating individuals, populations and species assemblages within ecosystems.

## Data Accessibility

All data used here are publicly available through two published data sets:

1. Raulo et al., 2022: Wild rodent trapping, tracking and sampling for genetics and gut microbiome, Silwood Park, Berkshire, November 2014 to November 2015. NERC EDS Environmental Information Data Centre. https://doi.org/10.5285/c67fde7f-a1c8-4cb4-a76e-0c6d21c82222
2. Raulo et al., 2023: Wild rodent tracking and gut microbiome data, Holly Hill, Wytham Woods, UK, 2018-2019. NERC EDS Environmental Information Data Centre. https://doi.org/10.5285/043513e5-406c-4477-89aa-c96059acb232

The code for deriving the social and spatial metrics and microbiome metrics used here, as well as the code for the models used in this article is available through Aura Raulo’s Git Hub page: https://github.com/nuorenarra/Interspecific-Microbiota-Transmission-

## Author contributions

AR led the data collection in Wytham, carried out all laboratory analyses, designed analytical methods, processed and analysed the data, and drafted the manuscript. SK led data collection from Silwood, planned laboratory analyses, designed analytical methods and provided feedback on the manuscript. JF helped design the methods for social network analysis and node metric calculation and provided feedback on the manuscript. EH helped develop the bioinformatics pipeline for the samples used here and provided feedback for the manuscript. CL led the development of tracking technology in Wytham and provided feedback on the manuscript.

## Funding Declaration

This work was carried out under the UK Home Office project license Animals (Scientific Procedures) Act (licence number PB0178858 held by S.C.L.K.). The research was supported by a grant from the National Centre for the Replacement, Refinement and Reduction of Animals in Research (NC3Rs; NC/R001103/1), a Natural Environment Research Council (NERC) fellowship (NE.L011867/1) to S.C.L.K., and funding from the European Research Council under the European Union’s Horizon 2020 research and innovation programme (grant agreement no. 851550). A.R. was supported by a Clarendon Scholarship, Kone Foundation and Queen’s College. J.A.F. acknowledges funding from the Biotechnology and Biological Sciences Research Council (BB/S009752/1) and NERC (NE/V013483/2).

## Acknowledgements

We wish to thank the wonderful research facilitators at Wytham and Silwood, in particular Nigel Fisher and the people who helped collect these unique data sets on rodent behaviour and gut microbiome: Bryony Allen, Saskia Ricks, Terence Chung, Selene Guitierrez Al-Khudhairy, Alice Weightman, Rohan Raval, Joe Williamson, Holly English and Kirsty Marsh. We thank Aurelio Malo for loan of PIT-tag loggers in Silwood and advice on field experimental design. We also want to thank Brendan Bohannan, Karen Guillemin and Laura Desban on insightful discussions over non-symmetrical transmission and Tim Coulson for guidance and feedback throughout this research project.

## Notes

https://doi.org/10.5285/c67fde7f-a1c8-4cb4-a76e-0c6d21c82222

https://doi.org/10.5285/043513e5-406c-4477-89aa-c96059acb232

https://github.com/nuorenarra/Interspecific-Microbiota-Transmission-

### Competing Interest Statement

The authors have declared no competing interest.

