## Supplementary material for "Interspecific social contacts spread gut microbiota in wild rodent communities"

**Index**

**Supplementary Appendices
Appendix S1:** Rationale behind the simplified spatial overlap metric

**Appendix S2:** Details on microbiota profiling with 16S rRNA marker gene

**Supplementary Figures**

**Figure S1.** The two types of RFID-loggers used at the two study sites. **Figure S2.** Temporal overlap of conspecific and heterospecific rodents in the tracking data.
**Figure S3**. Relationship between centroid distance and home range overlap
**Figure S4:** Distributions of social and spatial node metrics across species and sexes.
**Figure S5.** Pairplot of correlations between social and spatial connectedness measures

**Supplementary Tables
Table S1:** Additional information on the two study sites and data sets. **Table S2:** Results of Bayesian multi-membership betaregression models testing the effect of social association, spatial proximity and covariates on microbiota similarity (Jaccard Index) of conspecific (AS-AS) and heterospecific (AS-AF & AS-MG) rodent pairs in A) Wytham and B) Silwood.

**Table S3:** Results of Bayesian multi-membership betaregression models testing the effect of non-symmetrical social association on non-symmetrical microbiota similarity among AS-AF rodent pairs in Silwood, in the direction of A) AS-to-AF and B) AF-to-AS.

**Table S4:** Results of Bayesian Poisson regression models testing the effect of social and spatial connectedness to yellow-necked mice and conspecifics on AF-indicator taxa richness in wood mice in A) Wytham and B) Silwood.

**Table S5:** Results of Bayesian Poisson regression models testing for the effect of *extended* social connectedness to yellow-necked mice on AF-indicator taxa richness in wood mice with in A) Wytham and B) Silwood.

**Table S6:** Results of Bayesian Poisson regression models testing the effect of social and spatial connectedness to yellow-necked mice and conspecifics on complementary AS-indicator taxa richness in wood mice in A) Wytham and B) Silwood

**Table S7:** Results of Bayesian Poisson regression models testing the effect of social and spatial connectedness to bank voles and conspecifics on MG-indicator taxa richness in wood mice in Wytham

**Table S8:** Results of Bayesian Poisson regression models testing the effect of overall heterospecific and conspecific social and spatial connectedness on overall microbiota richness in wood mice in A) Wytham and B) Silwood

**Appendix S1. Home range analysis and the construction of simplified spatial proximity metrics**

Home ranges were constructed for Wytham rodents with denser logger data. Ranges were inferred as the 75% core of the autocorrelated kernel utilization density distribution ^1^, based on spatiotemporal (path-constructing) movement models implemented in R package *ctmm,* ^2^. For the home range analysis, logger data was first aggregated within ten-minute periods, such that all records of an individual at the same location within each ten minute-period were considered a single unique “logger visit”. To ensure that data quality was suitable for home range analysis we first filtered out all individuals that did not have at least five unique “logger visits” in at least three unique locations. We then inspected the remaining data using empirical variograms, a method visualising the autocorrelation structure in locational timeseries by presenting the semi-variance (proportional to home range size) as a function of increasing time lags between observations, with clear asymptotes diagnostic of range-residency^1^ . Based on this, we excluded individuals who’s variogram showed a positive linear relationship between estimated distance and time lag between logger observations, as these were considered signs of either logger histories that were too incomplete for reliable home range estimation (e.g., due to small amount of data or edge effects on the study grid) or spatially shifting home range location (unstable home range).

Using the data from the remaining Wytham rodent individuals we then fit a series of range-resident movement models using model selection to identify the best model fit, given the logger data^1^. We applied an autocorrelated kernel density estimator (AKDE) to these movement models using an error-informed model (with error set to be 1 meter, approximately the detection diameter of loggers). Although the observation model of fixed logger location data is different to the tracking data that the ADKE methods were developed for, the scales of animal movement observed here were larger than the grid resolution. Consequently, visual inspection of the kernels against the logging grid setup showed little evidence of excess density around the loggers. Once home ranges had been estimated, we calculated home range overlap for each rodent pair using the “overlap” function in *ctmm* and the Bhattacaryya coefficient, defined as:

$$BC(h,g)=\int\sqrt{h\left( x \right)g(x)} dx$$

, where *h* and *g* are two probability distributions (kernel density functions) of value *x* respectively.

Under these criteria, home ranges could be estimated for only a proportion of individuals in the Wytham data (52% of AF, 81% of AS, 73% of MG). Thus, we set out to develop a simpler measure of spatial proximity between individuals, that could be used with all individuals in the data (both Wytham and Silwood) and yet would be a good proxy of the home range overlap measured for this subset of Wytham individuals. Spatial proximity networks were constructed by first calculating the distances in meters between observation record centroids (mean locations) for each individual rodent. To assess whether this measure of centroid distance could provide a good proxy of home range overlap, and whether this applied up to a particular distance threshold, we examined the relationship between these two variables for the subset of Wytham rodents with dense enough tracking data for accurate home range estimates. For the 128 individuals (98 AS, 9 AF and 21 MG) from Wytham with sufficient home range estimates, we found that centroid distance (in meters) between pairs of (multiple species of) rodents was correlated with their Bhattacaryya home range overlap up to ~120 meters of centroid distance, after which the correlation broke down (Fig. S3). Based on this, for rodent pairs with centroid distance 0-120 m apart, we derived a measure of spatial proximity proximity by scaling these distances between 0 and 1 and taking the complement of the resulting value to transpose the metric into “proximity”. For rodent pairs with centroids >120 m apart, we set their spatial proximity to 0.

**Appendix S2.**

A) Microbiome profiling of Wytham samples:

*Sample analysis order*

The microbiome samples were analysed in two batches, each including i) DNA extraction, ii) library preparation and iii) sequencing on Illumina MiSeq platform. Both batches included soil samples as well as faecal samples of two rodent species (wood mice and yellow-necked mice). The first batch had additionally faecal samples from bank voles. All samples were randomised across 96-well plates for DNA extraction and re-ordered for 96-well plates for library preparation to enable detection of extraction and PCR-related technical effects on the final data. Sequencing batches had a slight temporal difference: First batch contained only samples from the first half of the study (February-July), while the second batch contained samples from throughout the study (February-November).

*DNA extraction*

DNA was extracted from samples with a 96-well plate format Zymo Quick-DNATM Fecal/Soil Microbe kits (Zymo) according to manufacturer's instructions, using a Qiagen TissueLyser plate-format bead beater for DNA homogenization. Each DNA extraction plate contained one mock community (ZYMOBiomics Microbial Community Standard cat. No. D6300) to evaluate extraction, PCR and sequencing accuracy. After extraction, DNA concentrations in samples and controls were quantified with a Qubit 3.0 Fluorometer to verify extraction success. Control samples were additionally amplified for bacterial DNA with a 40-cycle PCR (see below) and product run on gel to make sure no bacterial DNA amplified from negative controls. No bands were seen on these gels.

*Library preparation and sequencing*
Extracted DNA samples were re-ordered for library preparation plates and amplified in a two-step PCR with primers 515F and 926R targeting an approximately 370 bp sequence in the V4-V5 region of the bacterial 16S rRNA gene^3^ . Each plate contained 95 samples and one negative PCR-control. First round PCR had 20μl reaction volume, consisting of: 10μl KAPA 2x Mastermix (KAPA Biosystems), 0.25μl each primer at 10μM, 4.5μl ultra pure water and 5μl extracted DNA. Cycling conditions were as follows: denaturation at 98°C for 2min, 10 cycles of 95°C for 20s, 65°C for 15s, 70°C for 45s, followed by a final extension at 72°C for 5min and a 4°C hold. To reduce PCR- induced variation, this first round PCR was done in duplicates after which the products pooled (by mixing 10μl +10μl of each product on a clean plate) and purified on AMPure magnetic bead purification plate (according to manufacturer’s instructions). After purification, the second round PCR was prepared by mixing 9 μl of purified product with a 11 μl of master mix containing 10 μl KAPA Mastermix and 0.5+0.5 μl of forward and reverse Illumina Nextera barcoded indexing primers, added in unique combinations per each sample. The second PCR included 15 cycles with the same thermocycle conditions and was followed by the same AMPure purification step as the first PCR. Finally, amplified, indexed and purified 16S libraries were quantified for concentration (with Qubit), diluted to even concentration of 3-5 ng/ul and pooled for sequencing. Of each library preparation plate, the negative PCR control and 7 real samples (in diagonal order) were run on agarose gel in the end to ensure amplification success and lack of PCR-contamination. Extraction controls were not re-run on gels after amplification. Pooled libraries (x2) were sent off to the Centre for Genomic Research in Liverpool, where each pool was size-selected using a Pippin Prep, by excluding fragments outside the expected range. Libraries were then sequenced on two separate sequencing runs using 2x250bp paired-end sequencing on an Illumina MiSeq.

*Bioinformatics and preprocessing of microbiota sequence data*
Raw sequence data from each of the two sequencing batches was separately demultiplexed into their original samples and then processed through the DADA2 pipeline (version 1.14) ^4^. Here, *cutadapt^5^* was first used to determine optimal trimming length to remove primer and adapter sequences. Based on this, 24 base pairs from the beginning of each read were trimmed away using the trimLeft argument in FilterAndTrim function in DADA2. As part of this function, and following visual inspection of sequence quality, low- quality tails were also trimmed, leaving 290 bp for forward and 230 bp for reverse reads. Next, sequences were dereplicated (identical sequences combined and their quality scores merged), amplicon sequence variants (ASVs) were inferred using the DADA2 algorithm and reverse paired-end reads were merged. Reads that could not be merged or were of abnormal length (<365 bp or > 371 bp) were removed from the dataset. After this step, data from the two sequencing batches were combined and processed together: chimeric sequences were removed from the combined data and taxonomy was assigned to ASVs against the Silva Database ^6^ (version 138). Finally, the ASV-per-sample table, sample metadata table, ASV taxonomy table and phylogenetic tree of ASVs were combined into a “phyloseq object” using the R package *phyloseq^7^*. This phyloseq object is provided as part of this published data. Note that this phyloseq object is as raw as possible and has not gone through any of the customary preprocessing steps of microbiome data, such as dropping samples with low read count, filtering taxa based on prevalence or taxonomy, normalizing read counts etc.

We also want to note that we found evidence of contamination in some of our extraction plates: There were bacterial DNA in our control samples and in some plates, there was a spatial autocorrelation signal in microbiome similarity among samples extracted in the same plate, i.e., samples extracted closer to each other were more likely to share contaminant ASVs. Contamination was much stronger in extraction controls, including ones that went through a clean (uncontaminated) PCR, implying that our DNA extraction protocol was vulnerable to contamination. This is a common issue with plate-format extraction kits, such as the ZYMO 96-well plate format faecal/soil DNA extraction kit we used. This kind of contamination likely happens through spillage between samples during the lysis step, and is prone to increase “technical noise” in the microbiota data. While this noise is random across biological groups in our data (as our samples were randomised across extraction plates, it can potentially diminish or mask some real biological differences between samples. Because of this we decontaminated the phyloseq object before using it using *decontam* algorithm ^8^ with a probability threshold 0.75 to identify and filter out potential contaminants from this data. Here, to decontaminate the rodent samples, by using a phyloseq object containing only one sample type at a time (AS” or “AF” or “MG”) alongside water control samples and then running the decontam algorithm using this phyloseq object.

Visualization of mock community composition revealed that our laboratory- and bioinformatics pipeline successfully captured the microbial diversity present in mock samples, though relative abundances varied somewhat from their theoretically true levels

B) Microbiome profiling of Silwood samples:

*Library preparation and sequencing*

DNA was extracted from samples using Zymo Quick-DNATM Fecal/Soil Microbe 96 kits (Zymo) according to manufacturer's instructions. For extractions as well as subsequent PCRs, samples were randomized across batches/plates to avoid confounding biological and technical effects on downstream microbial data. An approximately 240bp V4 region of the 16S rRNA gene was amplified using primers N515F and N806R^9^ in two-step (tailed-tag) approach with dual- indexing^10^. In addition to samples, we included one extraction control (PCR- grade water subjected to the DNA extraction method) and one PCR control (PCR-grade water) per 96-well plate as well as 2 extractions of a mock community (ZYMOBiomics Microbial Community Standard cat. No. D6300) to evaluate extraction, PCR and sequencing accuracy.

For all samples a test first round PCR was performed, to check sample amplification and decide on an appropriate number of PCR cycles for the final library preparation. For this test PCR, the following were combined in a 20μl reaction volume: 10μl KAPA 2x Mastermix (KAPA Biosystems), 0.25μl each primer at 10μM, 4.5μl PCR-grade water and 5μl undiluted DNA extraction. Cycling conditions were as follows: denaturation at 98°C for 2min, 35 cycles of 95°C for 20s, 65°C for 15s, 70°C for 30s, followed by a final extension at 70°C for 5min and a 4°C hold. Products were visualized on 2% agarose gels, and no amplification was observed in extraction or PCR controls. Subsequently the first round PCR for use in subsequent sequencing was performed, using exactly the same conditions as described above but with fewer cycles (20 instead of 35).

Unpurified first round PCR products were sent to the Centre for Genomic Research (Liverpool, UK) for indexing PCRs, purification, pooling, amplicon size selection and sequencing. First round PCR products were purified using Agencourt Ampure XP beads (Beckman Coulter, Brea, CA, USA), by adding 20μl beads at room temperature to samples, before washing twice with 200μl 80% ethanol and eluting in 10μl H2O. For the second round PCR (addition of indices), the following were combined in a 20μl reaction volume: 10μl KAPA 2x Mastermix (KAPA Biosystems), 0.5μl of each primer at 10μM and 9μl of clean PCR product. For indexing, eight forward primers (i5) and twelve reverse (i7) primers were used, each containing a unique barcode, creating 96 unique combinations (barcode sequences are reported in the Illumina Nextera Protocol ^11^. Second round PCR products were purified using Agencourt Ampure XP beads as described above, but eluted in 20μl H2O. Each second round PCR product (library) was analyzed using a 2100 Bioanalyzer (Agilent Technologies, Santa Clara, CA, USA) and up to 96 libraries pooled per sequencing run, at equimolar concentration of fragments in the expected size range. Each pool was size-selected using a Pippin Prep, by excluding fragments outside the expected range. Libraries were sequenced using 2x250bp paired-end sequencing on an Illumina MiSeq, with samples from different extraction batches randomly allocated to four different runs. Visualization of the composition of the mock samples revealed that our laboratory- and bioinformatics pipeline successfully captured the microbial diversity present in mock samples

*Bioinformatics*

Sequence data were processed through the DADA2 pipeline (version 1.6.0) ^4^. *Cutadapt ^5^* command was used to determine optimal trimming parameters for removal of leading primer and adapter sequences (27 base pairs from the beginning of each read), and trimming was performed using the trimLeft argument in DADA2. Following visual inspection of sequence quality, low-quality tails were also trimmed (leaving 230 bp for forward and 180 bp for reverse reads). Sequences were dereplicated, and amplicon sequence variants (ASVs) inferred using the DADA2 algorithm. Forward and reverse paired-end reads were merged after which reads that could not be merged, were putative chimeric sequences or were of abnormal length (<238 bp or > 245 bp) were removed from the dataset.

Within the R package *phyloseq^7^* taxonomy was assigned to ASVs using the GreenGenes Database (GreenGenes Database Consortium 13.8). One sample with exceptionally low read count was removed from the dataset (n=300 reads, while read count >10 000 for all other samples). We then used package iNEXT^12^ to confirm that sample completeness estimates plateaued and diversity estimates stabilized at read counts around 2500-4000 reads, well below the read counts for all samples in the remaining dataset. Singleton reads were filtered out of the dataset, to avoid residual bias from possible contaminants or sequencing errors. ASVs assigned to taxa known to not be gut bacteria (Cyanobacteria, Xanthomonadales, Mitochondria) were also filtered out. Finally, ASV counts were normalized by calculating proportional abundance within each sample ^13^.

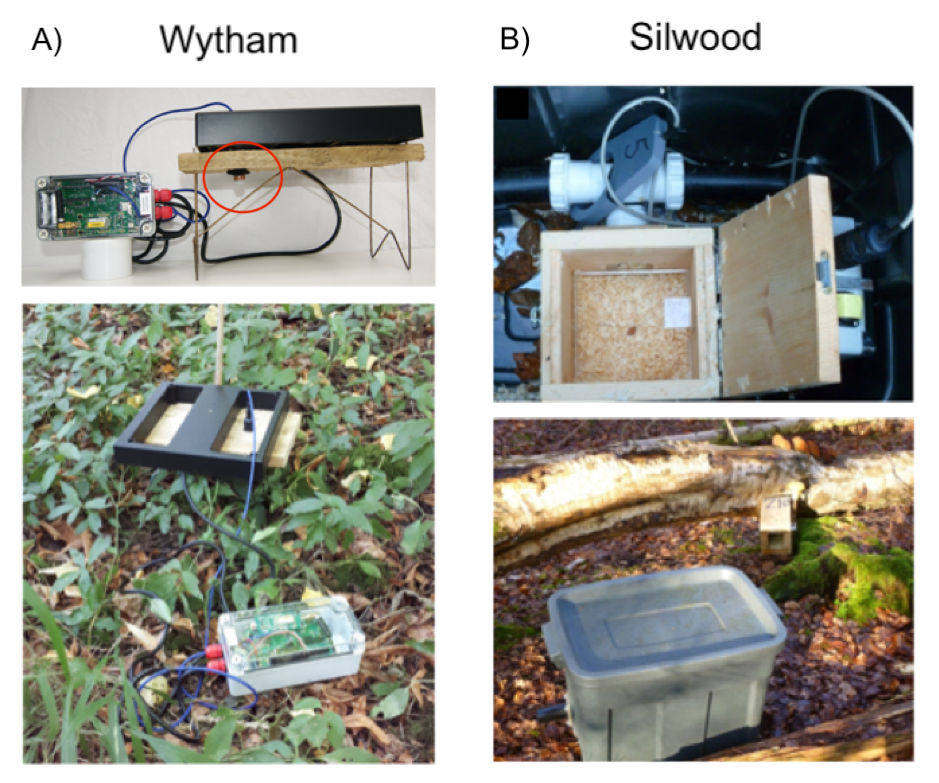

**Figure S1.** **The two types of RFID-loggers used in the two studies.**

A) In Wytham, the logger was unbaited and consisted of a motion sensor (red circle, figure top left), that woke up the detection coil (black square) whenever a warm object, such as a rodent came nearby. The coil, sitting on a wooden “table” to allow rodents to pass under as well as around it, then created a ~1m^2^ oscillating detection field that would interfere – and subsequently log - any 124 kHz RFID tag within this field. Exploiting this, wild rodents were tagged with a subcutaneous passive integrated transponder (PIT) tags containing a 10-digit HEX-code identification number readable in this exact oscillation frequency. When a rodent tag was read by the coil, this identification number was saved on an SD-card (in the grey box) and additionally transmitted into and interactive internet database through an antenna (in the grey box) connected to the broadband network covering Wytham Woods (Wytham Data Net).

B) In Silwood, the logger took the form of a large plastic box (figure bottom right) with two entrance tubes leading to a central wooden box containing sawdust and a single peanut (figure top right). An RFID reader coil surrounding the entrance to the central box was on continuously and recorded the PIT-tag ID of any tagged rodent present under it every 0.3 seconds. Peanut oil around the tube entrances and a single peanut inside the box, renewed at each logger rotation, was used as a minor lure. Upon each logger rotation, each logger box and entrance tubing were thoroughly cleaned with 70% ethanol and the sawdust was replaced. Cleaning was aided by the box’s removable plastic floor that could be cleaned thoroughly. Loggers were powered using 12V lead acid rechargeable batteries. Further details about these loggers can be found in ^14^.

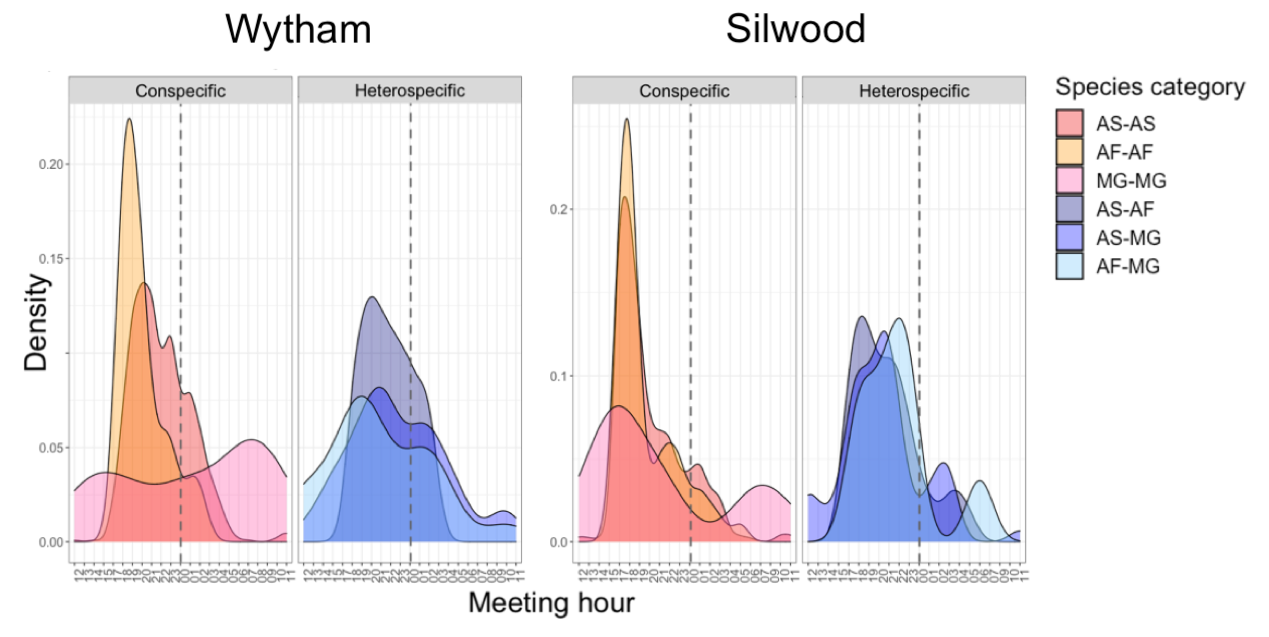

**Figure S2. Temporal overlap of conspecific and heterospecific rodents in the tracking data.** Distribution of co-occurrences by hour of day. Here, co-occurrences are defined as observations of two individuals at the same location (PIT-tag logger) within the same hour. Conspecific co-occurrences (warm colors) tend to happen during the general activity period of the species: During night hours for nocturnal mouse species (AS-AS, and AF-AF co-occurrences) and during light hours for more diurnal bank voles (MG-MG co-occurrences). Heterospecific co-occurrences (cold colors, AS-AF, AS MG, and AF-MG co-occurrences) tend to happen during the overlapping activity period of all three rodent species, a few hours before midnight (dashed line).

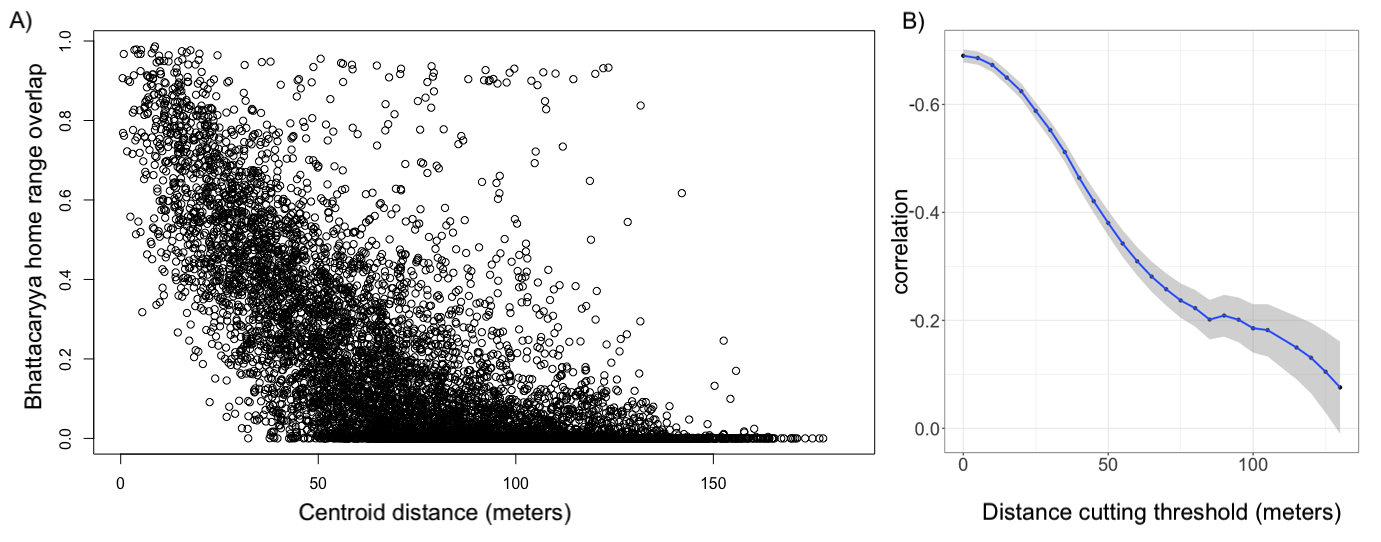

**Figure S3. Relationship between centroid distance and kernel density -based home range overlap**

A) Centroid distance is strongly negatively associated with home range overlap up to ~120 meters, but the correlation degrades thereafter. B) Considering increasingly faraway distances (x-axis, distance threshold below which rodent pairs were dropped from data), correlation (y-axis, Pearson’s correlation coefficient) decreases somewhat linearly. After 120 meters of centroid distance, the correlation (line) is not significant anymore (confidence intervals touch zero).

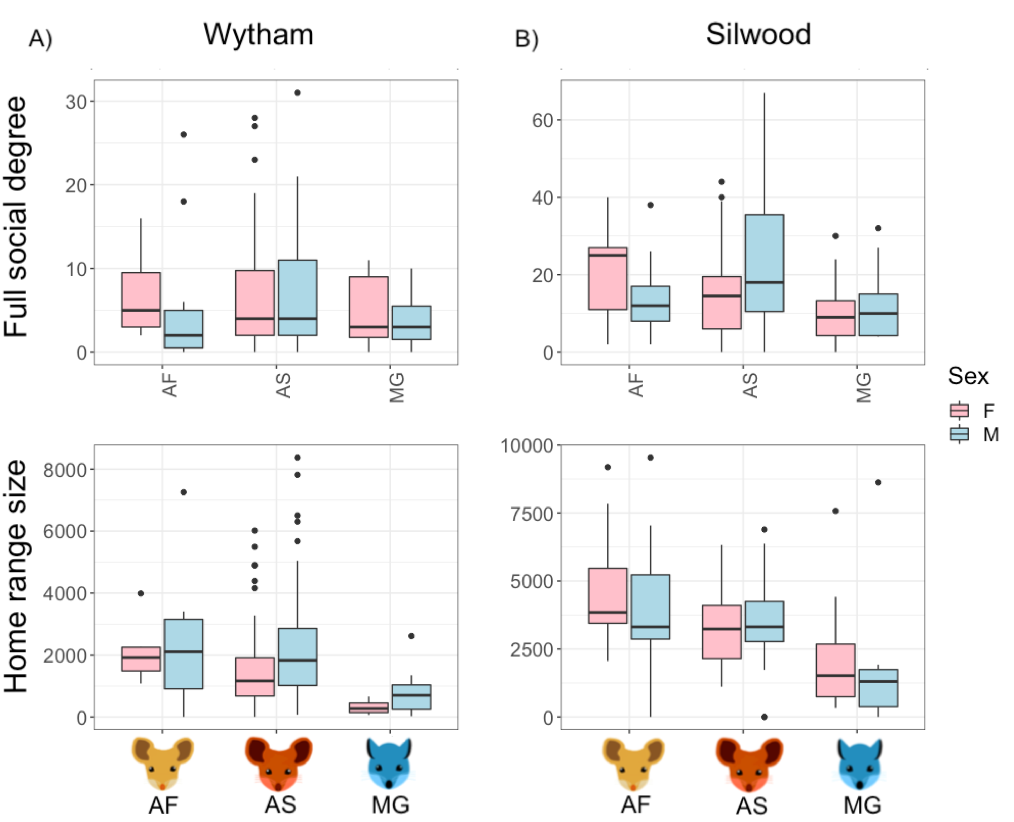

**Figure S4. Distributions of social and spatial node metrics across species and sexes.**

A) Full social degree (total number of direct social network connections to rodents of any species) had overlapping distributions across rodent species and in both Wytham and Silwood was highest in yellow-necked mouse females compared to other rodents. B) Home range sizes were greater for the two mouse species compared to voles and slightly larger for yellow-necked mice compared to wood mice. Yellow-necked mice had the largest home ranges (Wytham: mean 2199 m^2^, sd=1720 ; Silwood: mean 4321 m2, sd=2633), with wood mice having second largest (Wytham: mean 1909 m2, sd=1661; Silwood: mean 3367 m2, sd=1457), and bank voles having the smallest home ranges (Wytham: mean 550 m2, sd=550.1; Silwood: mean 1922 m2, sd=2047).

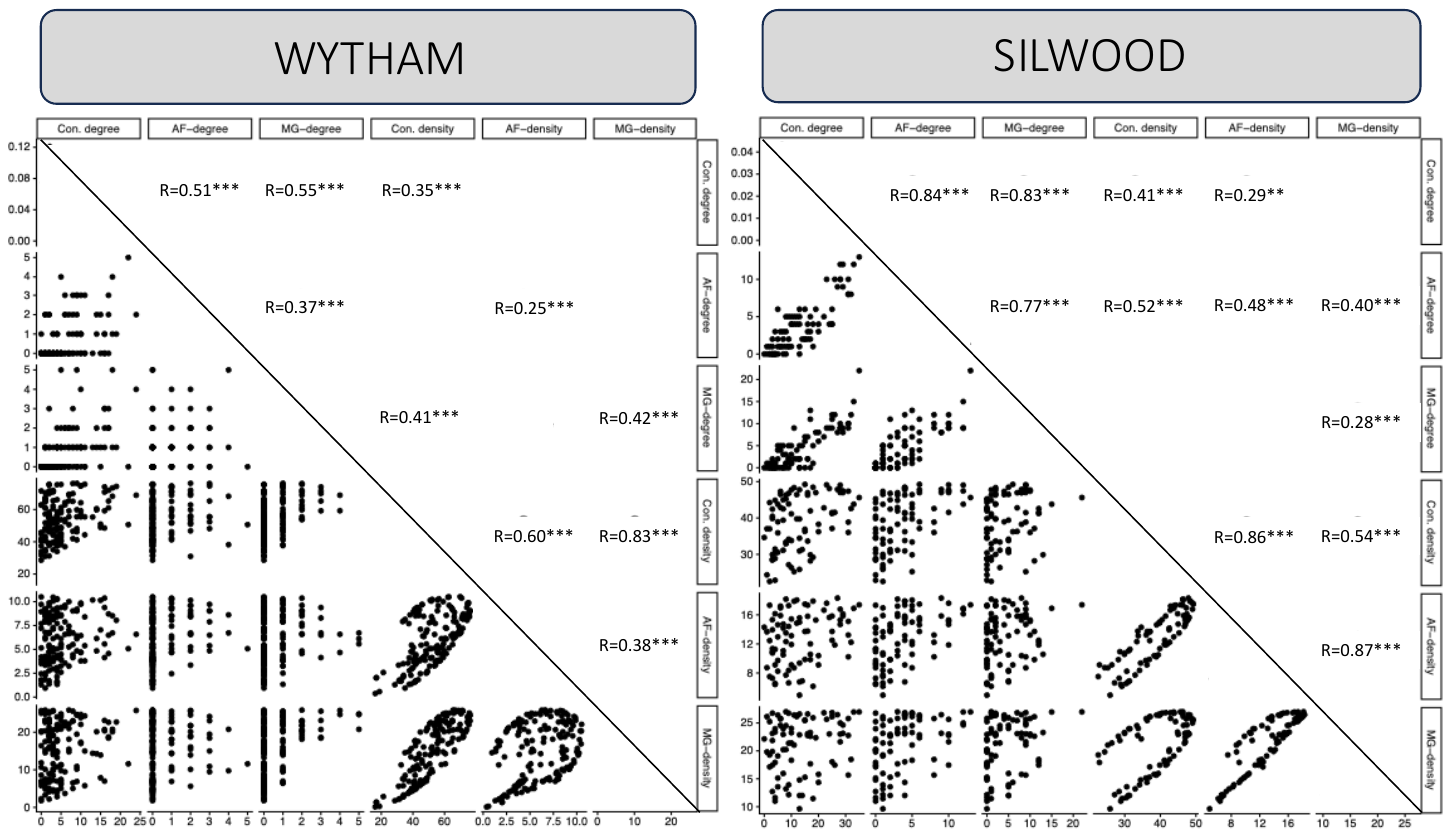

**Figure S5. Pairplot of correlations between social and spatial connectedness measures**
Covariation among connectedness metrics in A) Wytham data and B) Silwood data. Lower triangle visualizes correlation between intersecting variables and upper triangle reports significant pairwise correlations (Pearson’s correlation coefficient).

**Table S1**. **Additional information on the two study sites and data sets.**

Summarized differences between the (A) field data collection, (B) laboratory analyses and (C) resulting sample sizes in the two parallel studies.

| A) Field data collection: | | | | |
| --- | --- | --- | --- | --- |
|  | ***Wytham*** | | ***Silwood*** | |
| Study plot area | 2.56 ha | | 2.47 ha | |
| Study plot habitat type | open deciduous woodland, bramble bushes | | open deciduous woodland, rhododendron, bamboo | |
| Study duration and period | 10 months  (Feb-Nov 2019) | | 12 months  (Nov 2014-Nov 2015) | |
| Loggers used | 60 | | 9 | |
| Logger density | ~24.3 per ha | | ~3.5 per ha | |
| Logger design | - Large flat coil on a wooden table in open woodland. Coil reads PIT-tag of any rodent passing under or near the table within ~28cm - Loggers activated by body heat, when a warm object moves close to the PIR sensor   *See Figure S1A* | | - Plastic box with entrance tube. Circular coil around entrance tube, that reads the PIT-tag of any rodent that enters the chamber in the box centre - Loggers constantly on and recording data; Frequent battery recharging   *See Figure S1A* | |
| Logging design | - **Baiting**: no bait - **Logger placement and rotation**: Each logger had much smaller (400m^2^) territorty of 4 contiguous 10mx10m grid cells. Loggers were rotated fortnightly in a systematic design among the 4 grid cells, so that each logger covered its entire territory over a 2-month period. At any given moment, all loggers were positioned in an exactly even, chequerboard-like design across the site, with maximum distance to a logger being 15 m from any given point. | | - **Baiting**: Loggers baited with a single peanut in the inner chamber, and peanut oil around the rim of the entrance tube. - **Logger placement and rotation**: Each logger had a 2700 m^2^ territory of ~27 10x10m grid cells, within which it was rotated on average ~4 times per week. Upon rotation, a logger was positioned to a random cell within its territory (sampling cells without replacement) until the whole territory had been covered, before the process started again. Details can be found from ^15^ | |
| B) Laboratory analyses | | | | |
|  | ***Wytham*** | | ***Silwood*** | |
| Microbiota sample DNA extraction | Using Zymo fecal/Soil DNA plate format DNA extraction kit with Qiagen Tissuelyzer II | | Using Zymo fecal/Soil plate-format DNA extraction kit with a tube format Tissulyzer | |
| Microbiota sample Library preparation | Using 16S V4-5 primers 515F and 926R^3^ and a two-step (tailed-tag) approach with dual-indexing. Details in Appendix S2 | | Using 16S V4 primers 515F and 805R^9^ and a two-step (tailed-tag) approach with dual-indexing. Details in Appendix S2 | |
| Bioinformatics | Sample preprocessing done with DADA2 pipeline (1.14.0) Taxonomy assigned against Silva data base. Details in Appendix S2 | | Sample preprocessing done with DADA2 pipeline (1.6.0) Taxonomy assigned against GreenGenes Database (GreenGenes Consortium 13.8). Details in Appendix S2 | |
| C) Sample size summary | | | | |
|  | ***Wytham*** | | ***Silwood*** | |
|  | Individuals recorded on loggers | Microbiota samples from these individuals | Individuals recorded on loggers | Microbiota samples from these individuals |
| Wood mice (AS) | 150 | 297 | 83 | 239 |
| Yellow-necked mice (AF) | 18 | 21 | 26 | 30 |
| Bank voles (MG) | 35 | 22 | 49 | 0 |

**Table S2: Results of Bayesian multi-membership betaregression models testing the effect of social association, spatial proximity and covariates on microbiota similarity (Jaccard Index) of conspecific (AS-AS) and heterospecific (AS-AF & AS-MG) rodent pairs in A) Wytham and B) Silwood.**

Main predictors of interest are shown in bold, while control variables are shown in light grey. Values of all predictors and covariates ranged between 0-1 or were scaled to do so. Significant terms (where 95% credible intervals do not overlap zero) are shown in red. Post.mean indicates the mean of the posterior distribution of effect estimates, while lCI and uCI show the lower and upper limits of 95% credible intervals respectively. Est.Error indicates the standard deviation of the posterior distribution. For ease of interpretation, interaction effects are *NOT* reported as slopes relative to each other but as independent effects within each group.

**Table S2A**

Model formula: Jaccard microbiota similarity ~Read_depth_difference + Pcr_similarity + Extraction_similarity + Sex_similarity + Age_similarity + Sampling_interval + Species_pair + Species_pair*Social_association + Species_pair*Spatial_proximity + (1|mm(Individual_A,Individual_B)) + (1|mm(Sample_A,Sample_B))

| **Predictor** | **Estimate** | **lCI** | **uCI** | **Est.error** |
| --- | --- | --- | --- | --- |
| (Intercept) | -0.98 | -1.04 | -0.92 | 0.029 |
| Read depth difference | -0.24 | -0.26 | -0.23 | 0.008 |
| Pcr similarity | 0.03 | 0.03 | 0.04 | 0.002 |
| Extraction similarity | 0.00 | 0.00 | 0.01 | 0.002 |
| Sex similarity | 0.00 | 0.00 | 0.00 | 0.002 |
| Age similarity | 0.12 | 0.09 | 0.15 | 0.014 |
| Sampling interval | -0.28 | -0.29 | -0.27 | 0.004 |
| Species pair:AF-AS | -0.35 | -0.45 | -0.26 | 0.050 |
| Species pair:AS-MG | -2.35 | -2.46 | -2.25 | 0.054 |
| **AS-AS social association** | **0.26** | **0.21** | **0.31** | **0.025** |
| **AS-AF social association** | **0.03** | **-0.16** | **0.20** | **0.099** |
| **AS-MG social association** | **0.10** | **-0.63** | **0.86** | **0.387** |
| **AS-AS spatial proximity** | **0.06** | **0.05** | **0.06** | **0.003** |
| **AS-AF spatial proximity** | **0.05** | **0.03** | **0.06** | **0.010** |
| **AS-MG spatial proximity** | **-0.04** | **-0.08** | **0.01** | **0.022** |

**Table S2B**

Model formula: Jaccard microbiota similarity ~Read_depth_difference + Pcr_similarity + Sex_similarity + Age_similarity + Sampling_interval + Species_pair + Species_pair*Social_association + Species_pair*Spatial_proximity + (1|mm(Individual_A,Individual_B)) + (1|mm(Sample_A,Sample_B))

| **Predictor** | **Estimate** | **lCI** | **uCI** | **Est.error** |
| --- | --- | --- | --- | --- |
| (Intercept) | -1.22 | -1.29 | -1.15 | 0.036 |
| Read depth difference | -0.04 | -0.15 | 0.07 | 0.054 |
| Pcr similarity | 0.01 | 0.00 | 0.01 | 0.002 |
| Sex similarity | 0.00 | 0.00 | 0.01 | 0.002 |
| Age similarity | -0.01 | -0.04 | 0.02 | 0.018 |
| Sampling interval | -0.51 | -0.52 | -0.50 | 0.005 |
| Species pair:AF-AS | -0.03 | -0.12 | 0.05 | 0.045 |
| **AS-AS social association** | **0.52** | **0.48** | **0.56** | **0.021** |
| **AS-AF social association** | **0.27** | **0.20** | **0.35** | **0.046** |
| **AS-AS spatial proximity** | **0.03** | **0.02** | **0.04** | **0.006** |
| **AS-AF spatial proximity** | **0.04** | **0.02** | **0.06** | **0.013** |

**Table S3: Results of Bayesian multi-membership betaregression models testing the effect of non-symmetrical social association on non-symmetrical microbiota similarity among AS-AF rodent pairs in Silwood, in the direction of A) AS-to-AF and B) AF-to-AS.**

Main predictors of interest are shown in bold, while control variables are shown in light grey. Values of all predictors and covariates ranged between 0-1 or were scaled to do so. Significant terms (where 95% credible intervals do not overlap zero) are shown in red. Post.mean indicates the mean of the posterior distribution of effect estimates, while lCI and uCI show the lower and upper limits of 95% credible intervals respectively. Est.Error indicates the standard deviation of the posterior distribution.

**Table S3A**

Model formula: Non-symmetrical Jaccard microbiota similarity ~Read_depth_difference + Pcr_similarity + Sex_similarity + Age_similarity + Sampling_interval + Spatial_proximity +
AS-to-AF_social_association + (1|mm(Individual_A,Individual_B)) + (1|mm(Sample_A,Sample_B))

| **Predictor** | **Estimate** | **lCI** | **uCI** | **Est.error** |
| --- | --- | --- | --- | --- |
| (Intercept) | -0.83 | -1.01 | -0.65 | 0.09 |
| Read depth difference | -0.03 | -0.08 | 0.02 | 0.02 |
| Pcr similarity | 0.00 | -0.01 | 0.01 | 0.01 |
| Sex similarity | -0.00 | -0.01 | 0.01 | 0.01 |
| Age similarity | 0.25 | 0.09 | 0.41 | 0.08 |
| Sampling interval | -0.31 | -0.35 | -0.28 | 0.02 |
| Spatial proximity | 0.00 | -0.04 | 0.05 | 0.02 |
| **AS-to-AF social association** | **0.10** | **0.05** | **0.16** | **0.03** |

**Table S3B**

Model formula: Non-symmetrical Jaccard microbiota similarity ~Read_depth_difference + Pcr_similarity + Sex_similarity + Age_similarity + Sampling_interval + Spatial_proximity +
AF-to-AS_social_association + (1|mm(Individual_A,Individual_B)) + (1|mm(Sample_A,Sample_B))

| **Predictor** | **Estimate** | **lCI** | **uCI** | **Est.error** |
| --- | --- | --- | --- | --- |
| (Intercept) | -0.57 | -0.83 | -0.32 | 0.13 |
| Read depth difference | -0.06 | -0.10 | -0.01 | 0.03 |
| Pcr similarity | 0.00 | -0.01 | 0.02 | 0.01 |
| Sex similarity | -0.01 | -0.02 | 0.01 | 0.01 |
| Age similarity | 0.10 | -0.12 | 0.32 | 0.11 |
| Sampling interval | -0.33 | -0.37 | -0.29 | 0.02 |
| Spatial proximity | -0.01 | -0.06 | 0.04 | 0.03 |
| **AF-to-AS social association** | **0.27** | **0.21** | **0.33** | **0.03** |

**Table S4: Results of Bayesian Poisson regression models testing the effect of social and spatial connectedness to yellow-necked mice and conspecifics on AF-indicator taxa richness in wood mice in A) Wytham and B) Silwood.**

Main predictors of interest are shown in bold, while control variables are shown in light grey. Values of all predictors and covariates ranged between 0-1 or were scaled to do so. Significant terms (where 95% credible intervals do not overlap zero) are shown in red. Post.mean indicates the mean of the posterior distribution of effect estimates, while lCI and uCI show the lower and upper limits of 95% credible intervals respectively. Est.Error indicates the standard deviation of the posterior distribution.

**Table S4A**

Model formula: AF-taxa_richness ~Read depth + Sex + Month + Age + Microbiota richness +
Conspecific density + Conspecific social degree + AF density + AF social degree + 1|Individual ID + 1| Pcr plate + 1| Extraction batch + 1| Sequencing batch

| **Predictor** | **post.mean** | **lCI** | **uCI** | **Est.error** |
| --- | --- | --- | --- | --- |
| (Intercept) | 2.51 | 2.26 | 2.74 | 0.55 |
| Read Depth | 0.34 | 0.19 | 0.49 | 0.09 |
| Sex:M | 0.02 | -0.03 | 0.07 | 0.03 |
| month:18-12 | -0.40 | -0.86 | 0.05 | 0.28 |
| month:19-01 | 0.29 | 0.08 | 0.51 | 0.11 |
| month:19-02 | 0.21 | 0.10 | 0.35 | 0.07 |
| month:19-03 | 0.34 | 0.23 | 0.51 | 0.07 |
| month:19-04 | 0.49 | 0.36 | 0.68 | 0.09 |
| month:19-05 | 0.45 | 0.15 | 0.70 | 0.14 |
| month:19-06 | 0.29 | 0.13 | 0.46 | 0.09 |
| month:19-07 | 0.36 | 0.25 | 0.51 | 0.07 |
| month:19-08 | 0.42 | 0.27 | 0.59 | 0.09 |
| month:19-09 | 0.34 | 0.20 | 0.51 | 0.09 |
| month:19-10 | 0.31 | 0.15 | 0.47 | 0.09 |
| month:19-11 | 0.38 | 0.19 | 0.56 | 0.10 |
| Age:Juvenile | -0.22 | -0.36 | -0.11 | 0.06 |
| Microbiome Richness | 0.55 | 0.35 | 0.77 | 0.12 |
| **Conspecific density** | **0.03** | **-0.12** | **0.15** | **0.07** |
| **Conspecific social degree** | **0.09** | **-0.05** | **0.22** | **0.07** |
| **AF density** | **-0.03** | **-0.15** | **0.09** | **0.06** |
| **AF social degree** | **0.19** | **0.07** | **0.33** | **0.07** |

**Table S4B**

Model formula: AF-taxa_richness ~Read_depth + Sex + Month + Age + Microbiota richness +
Conspecific density + Conspecific social degree + AF density + AF social degree + 1|Individual_ID + 1| Pcr_plate

| **Predictor** | **post.mean** | **lCI** | **uCI** | **Est.error** |
| --- | --- | --- | --- | --- |
| (Intercept) | 3.43 | 3.38 | 3.51 | 0.03 |
| Read Depth | -0.16 | -0.21 | -0.05 | 0.04 |
| Sex:M | -0.05 | -0.07 | 0.00 | 0.02 |
| month:02 | 0.06 | -0.02 | 0.11 | 0.03 |
| month:03 | 0.05 | -0.01 | 0.08 | 0.03 |
| month:04 | 0.09 | 0.02 | 0.13 | 0.03 |
| month:05 | 0.01 | -0.04 | 0.05 | 0.02 |
| month:06 | 0.00 | -0.09 | 0.04 | 0.04 |
| month:07 | -0.13 | -0.21 | 0.05 | 0.06 |
| month:08 | -0.03 | -0.08 | 0.02 | 0.03 |
| month:09 | -0.03 | -0.15 | 0.03 | 0.06 |
| month:10 | -0.04 | -0.15 | 0.02 | 0.05 |
| month:11 | -0.03 | -0.08 | 0.01 | 0.02 |
| month:12 | -0.01 | -0.12 | 0.05 | 0.04 |
| Age:Juvenile | -0.21 | -0.27 | -0.13 | 0.04 |
| Microbiome Richness | 0.94 | 0.83 | 1.01 | 0.05 |
| **Conspecific density** | **-0.08** | **-0.23** | **0.01** | **0.06** |
| **Conspecific social degree** | **-0.07** | **-0.20** | **0.01** | **0.06** |
| **AF density** | **0.04** | **-0.04** | **0.17** | **0.05** |
| **AF social degree** | **0.19** | **0.14** | **0.36** | **0.05** |

**Table S5: Results of Bayesian Poisson regression models testing for the effect of *extended* social connectedness to yellow-necked mice on AF-indicator taxa richness in wood mice with in A) Wytham and B) Silwood.**

Main predictors of interest are shown in bold, while control variables are shown in light grey. Values of all predictors and covariates ranged between 0-1 or were scaled to do so. Significant terms (where 95% credible intervals do not overlap zero) are shown in red. Post.mean indicates the mean of the posterior distribution of effect estimates, while lCI and uCI show the lower and upper limits of 95% credible intervals respectively. Est.Error indicates the standard deviation of the posterior distribution.

**Table S5A**

Model formula: AF-taxa_richness ~Read depth + Sex + Month + Age + Microbiota richness +
Conspecific density + Conspecific social degree + AF density + Extended AF social degree + 1|Individual ID + 1| Pcr plate + 1| Extraction batch + 1| Sequencing batch

| **Predictor** | **post.mean** | **lCI** | **uCI** | **Est.error** |
| --- | --- | --- | --- | --- |
| (Intercept) | 2.57 | 2.29 | 2.73 | 1.06 |
| Read Depth | 0.25 | 0.05 | 0.39 | 0.11 |
| Sex:M | 0.01 | -0.05 | 0.06 | 0.03 |
| month:18-12 | -0.23 | -0.57 | 0.10 | 0.23 |
| month:19-01 | 0.31 | 0.16 | 0.45 | 0.08 |
| month:19-02 | 0.14 | 0.03 | 0.25 | 0.06 |
| month:19-03 | 0.30 | 0.16 | 0.40 | 0.06 |
| month:19-04 | 0.44 | 0.32 | 0.58 | 0.07 |
| month:19-05 | 0.43 | 0.30 | 0.56 | 0.08 |
| month:19-06 | 0.22 | 0.04 | 0.37 | 0.10 |
| month:19-07 | 0.32 | 0.21 | 0.46 | 0.07 |
| month:19-08 | 0.35 | 0.25 | 0.57 | 0.08 |
| month:19-09 | 0.30 | 0.20 | 0.49 | 0.07 |
| month:19-10 | 0.23 | 0.14 | 0.40 | 0.07 |
| month:19-11 | 0.35 | 0.24 | 0.55 | 0.09 |
| Age:Juvenile | -0.19 | -0.37 | -0.04 | 0.09 |
| Microbiome Richness | 0.60 | 0.47 | 0.72 | 0.08 |
| **Conspecific density** | **0.03** | **-0.11** | **0.16** | **0.07** |
| **Conspecific social degree** | **0.07** | **-0.07** | **0.20** | **0.07** |
| **AF density** | **0.04** | **-0.05** | **0.13** | **0.05** |
| **Extended AF social degree** | **0.18** | **0.05** | **0.31** | **0.07** |

**Table S5B**

Model formula: AF-taxa_richness ~Read depth + Sex + Month + Age + Microbiota richness +
Conspecific density + Conspecific social degree + AF density + Extended AF social degree + 1|Individual ID + 1| Pcr plate

| **Predictor** | **post.mean** | **lCI** | **uCI** | **Est.error** |
| --- | --- | --- | --- | --- |
| (Intercept) | 2.57 | 2.29 | 2.73 | 1.06 |
| Read Depth | 0.25 | 0.05 | 0.39 | 0.11 |
| Sex:M | 0.01 | -0.05 | 0.06 | 0.03 |
| month:18-12 | -0.23 | -0.57 | 0.10 | 0.23 |
| month:19-01 | 0.31 | 0.16 | 0.45 | 0.08 |
| month:19-02 | 0.14 | 0.03 | 0.25 | 0.06 |
| month:19-03 | 0.30 | 0.16 | 0.40 | 0.06 |
| month:19-04 | 0.44 | 0.32 | 0.58 | 0.07 |
| month:19-05 | 0.43 | 0.30 | 0.56 | 0.08 |
| month:19-06 | 0.22 | 0.04 | 0.37 | 0.10 |
| month:19-07 | 0.32 | 0.21 | 0.46 | 0.07 |
| month:19-08 | 0.35 | 0.25 | 0.57 | 0.08 |
| month:19-09 | 0.30 | 0.20 | 0.49 | 0.07 |
| month:19-10 | 0.23 | 0.14 | 0.40 | 0.07 |
| month:19-11 | 0.35 | 0.24 | 0.55 | 0.09 |
| Age:Juvenile | -0.19 | -0.37 | -0.04 | 0.09 |
| Microbiome Richness | 0.60 | 0.47 | 0.72 | 0.08 |
| **Conspecific density** | **0.03** | **-0.11** | **0.16** | **0.07** |
| **Conspecific social degree** | **0.07** | **-0.07** | **0.20** | **0.07** |
| **AF density** | **0.04** | **-0.05** | **0.13** | **0.05** |
| **Extended AF social degree** | **0.18** | **0.05** | **0.31** | **0.07** |

**Table S6: Results of Bayesian Poisson regression models testing the effect of social and spatial connectedness to yellow-necked mice and conspecifics on complementary AS-indicator taxa richness in wood mice in A) Wytham and B) Silwood.**

Main predictors of interest are shown in bold, while control variables are shown in light grey. Values of all predictors and covariates ranged between 0-1 or were scaled to do so. Significant terms (where 95% credible intervals do not overlap zero) are shown in red. Post.mean indicates the mean of the posterior distribution of effect estimates, while lCI and uCI show the lower and upper limits of 95% credible intervals respectively. Est.Error indicates the standard deviation of the posterior distribution.

**Table S6A**

Model formula: AS-taxa_richness ~Read depth + Sex + Month + Age + Microbiota richness +
Conspecific density + Conspecific social degree + AF density + AF social degree + 1|Individual ID + 1| Pcr plate + 1| Extraction batch + 1| Sequencing batch

| **Predictor** | **post.mean** | **lCI** | **uCI** | **Est.error** |
| --- | --- | --- | --- | --- |
| (Intercept) | 3.83 | 3.75 | 3.91 | 0.10 |
| Read Depth | 0.00 | -0.09 | 0.11 | 0.06 |
| Sex:M | -0.02 | -0.05 | 0.01 | 0.01 |
| month:18-12 | 0.01 | -0.24 | 0.21 | 0.12 |
| month:19-01 | -0.01 | -0.15 | 0.18 | 0.10 |
| month:19-02 | 0.01 | -0.05 | 0.05 | 0.03 |
| month:19-03 | 0.03 | -0.04 | 0.10 | 0.04 |
| month:19-04 | -0.01 | -0.08 | 0.06 | 0.04 |
| month:19-05 | -0.06 | -0.19 | 0.05 | 0.06 |
| month:19-06 | 0.07 | -0.02 | 0.13 | 0.03 |
| month:19-07 | 0.00 | -0.08 | 0.08 | 0.04 |
| month:19-08 | -0.01 | -0.08 | 0.07 | 0.04 |
| month:19-09 | 0.00 | -0.08 | 0.09 | 0.05 |
| month:19-10 | -0.01 | -0.07 | 0.04 | 0.03 |
| month:19-11 | 0.02 | -0.05 | 0.12 | 0.05 |
| Age:Juvenile | -0.01 | -0.09 | 0.07 | 0.05 |
| Microbiome Richness | 0.96 | 0.86 | 1.03 | 0.05 |
| **Conspecific density** | **0.02** | **-0.05** | **0.09** | **0.03** |
| **Conspecific social degree** | **0.05** | **-0.01** | **0.12** | **0.04** |
| **AF density** | **0.00** | **-0.07** | **0.06** | **0.03** |
| **AF social degree** | **-0.07** | **-0.12** | **-0.01** | **0.03** |

**Table S6B**

Model formula: AS-taxa_richness ~Read depth + Sex + Month + Age + Microbiota richness +
Conspecific density + Conspecific social degree + AF density + AF social degree + 1|Individual ID + 1| Pcr plate

| **Predictor** | **post.mean** | **lCI** | **uCI** | **Est.error** |
| --- | --- | --- | --- | --- |
| (Intercept) | 3.55 | 3.50 | 3.65 | 0.05 |
| Read Depth | -0.08 | -0.16 | 0.01 | 0.04 |
| Sex:M | 0.05 | 0.02 | 0.07 | 0.01 |
| month:02 | 0.04 | 0.00 | 0.12 | 0.03 |
| month:03 | 0.03 | -0.01 | 0.08 | 0.02 |
| month:04 | 0.02 | -0.05 | 0.07 | 0.03 |
| month:05 | 0.02 | -0.10 | 0.10 | 0.07 |
| month:06 | -0.05 | -0.12 | 0.00 | 0.04 |
| month:07 | 0.12 | -0.02 | 0.18 | 0.06 |
| month:08 | -0.03 | -0.08 | 0.02 | 0.02 |
| month:09 | -0.02 | -0.13 | 0.03 | 0.04 |
| month:10 | -0.11 | -0.19 | -0.04 | 0.04 |
| month:11 | 0.02 | -0.04 | 0.05 | 0.03 |
| month:12 | -0.01 | -0.05 | 0.05 | 0.03 |
| Age:Juvenile | 0.04 | -0.02 | 0.10 | 0.03 |
| Microbiome Richness | 1.07 | 0.98 | 1.12 | 0.03 |
| **Conspecific density** | **-0.11** | **-0.19** | **0.03** | **0.05** |
| **Conspecific social degree** | **0.14** | **0.09** | **0.22** | **0.03** |
| **AF density** | **0.10** | **-0.07** | **0.19** | **0.07** |
| **AF social degree** | **-0.12** | **-0.18** | **-0.05** | **0.03** |

**Table S7: Results of Bayesian Poisson regression models testing the effect of social and spatial connectedness to bank voles and conspecifics on complementary MG-indicator taxa richness in wood mice in Wytham**

Main predictors of interest are shown in bold, while control variables are shown in light grey. Values of all predictors and covariates ranged between 0-1 or were scaled to do so. Significant terms (where 95% credible intervals do not overlap zero) are shown in red. Post.mean indicates the mean of the posterior distribution of effect estimates, while lCI and uCI show the lower and upper limits of 95% credible intervals respectively. Est.Error indicates the standard deviation of the posterior distribution.

**Table S7**

Model formula: MG-taxa richness ~Read depth + Sex + Month + Age + Microbiota richness +
Conspecific density + Conspecific social degree + AF density + AF social degree + 1|Individual ID + 1| Pcr plate + 1| Extraction batch + 1| Sequencing batch

| **Predictor** | **post.mean** | **lCI** | **uCI** | **Est.error** |
| --- | --- | --- | --- | --- |
| (Intercept) | -0.56 | -1.67 | 0.03 | 0.46 |
| Read Depth | -1.07 | -1.43 | -0.07 | 0.40 |
| Sex:M | -0.10 | -0.24 | 0.09 | 0.07 |
| month:18-12 | -0.99 | -1.52 | 1.61 | 0.83 |
| month:19-01 | -0.09 | -0.63 | 2.27 | 0.80 |
| month:19-02 | 0.29 | -0.08 | 0.87 | 0.21 |
| month:19-03 | 0.69 | 0.11 | 1.22 | 0.29 |
| month:19-04 | 0.63 | 0.11 | 1.50 | 0.34 |
| month:19-05 | 0.80 | 0.02 | 2.29 | 0.53 |
| month:19-06 | 0.33 | -0.22 | 0.95 | 0.37 |
| month:19-07 | 0.35 | -0.06 | 1.34 | 0.31 |
| month:19-08 | 0.47 | 0.03 | 1.25 | 0.30 |
| month:19-09 | -0.84 | -1.13 | 0.52 | 0.42 |
| month:19-10 | 0.41 | -0.07 | 1.24 | 0.30 |
| month:19-11 | -0.21 | -0.68 | 0.65 | 0.35 |
| Age:Juvenile | 0.86 | 0.13 | 1.17 | 0.32 |
| Microbiome Richness | 1.09 | 0.54 | 1.44 | 0.18 |
| **Conspecific density** | **-0.25** | **-0.97** | **0.15** | **0.30** |
| **Conspecific social degree** | **0.20** | **-0.37** | **0.62** | **0.24** |
| **MG density** | **0.47** | **0.07** | **1.28** | **0.30** |
| **MG social degree** | **-0.57** | **-0.91** | **0.04** | **0.25** |

**Table S8: Results of Bayesian Poisson regression models testing the effect of overall heterospecific and conspecific social and spatial connectedness on overall microbiota richness in wood mice in A) Wytham and B) Silwood**

Main predictors of interest are shown in bold, while control variables are shown in light grey. Values of all predictors and covariates ranged between 0-1 or were scaled to do so. Significant terms (where 95% credible intervals do not overlap zero) are shown in red. Post.mean indicates the mean of the posterior distribution of effect estimates, while lCI and uCI show the lower and upper limits of 95% credible intervals respectively. Est.Error indicates the standard deviation of the posterior distribution.

**Table S8A**

Model formula: Microbiota richness ~Read depth + Sex + Month + Age +
Conspecific density + Conspecific social degree + Heterospecific density + Heterospecific social degree + 1|Individual ID + 1| Pcr plate + 1| Extraction batch + 1| Sequencing batch

| **Predictor** | **post.mean** | **lCI** | **uCI** | **Est.error** |
| --- | --- | --- | --- | --- |
| (Intercept) | 5.09 | 4.89 | 5.29 | 0.26 |
| Read Depth | 0.74 | 0.53 | 0.96 | 0.11 |
| Sex:M | 0.00 | -0.05 | 0.05 | 0.03 |
| month:18-12 | -0.19 | -0.54 | 0.15 | 0.18 |
| month:19-01 | -0.12 | -0.39 | 0.20 | 0.15 |
| month:19-02 | 0.03 | -0.12 | 0.19 | 0.08 |
| month:19-03 | 0.01 | -0.16 | 0.17 | 0.08 |
| month:19-04 | -0.03 | -0.21 | 0.12 | 0.09 |
| month:19-05 | 0.19 | -0.16 | 0.52 | 0.17 |
| month:19-06 | 0.07 | -0.14 | 0.25 | 0.10 |
| month:19-07 | 0.09 | -0.08 | 0.28 | 0.09 |
| month:19-08 | -0.02 | -0.19 | 0.16 | 0.09 |
| month:19-09 | 0.00 | -0.17 | 0.19 | 0.09 |
| month:19-10 | -0.13 | -0.28 | 0.06 | 0.09 |
| month:19-11 | -0.06 | -0.26 | 0.12 | 0.09 |
| Age:Juvenile | -0.06 | -0.22 | 0.10 | 0.08 |
| **Conspecific density** | **0.11** | **-0.10** | **0.33** | **0.11** |
| **Conspecific social degree** | **0.05** | **-0.12** | **0.21** | **0.09** |
| **Heterospecific density** | **-0.15** | **-0.36** | **0.03** | **0.10** |
| **Heterospecific social degree** | **0.03** | **-0.13** | **0.21** | **0.09** |

**Table S8B**

Model formula: Microbiota richness ~Read depth + Sex + Month + Age +
Conspecific density + Conspecific social degree + Heterospecific density + Heterospecific social degree + 1|Individual ID + 1| Pcr plate

| **Predictor** | **post.mean** | **lCI** | **uCI** | **Est.error** |
| --- | --- | --- | --- | --- |
| (Intercept) | 5.09 | 4.87 | 5.27 | 0.10 |
| Read Depth | 0.48 | 0.29 | 0.70 | 0.11 |
| Sex:M | 0.04 | -0.04 | 0.11 | 0.04 |
| month:02 | 0.17 | 0.01 | 0.31 | 0.08 |
| month:03 | 0.15 | 0.01 | 0.28 | 0.07 |
| month:04 | 0.10 | -0.06 | 0.28 | 0.09 |
| month:05 | 0.04 | -0.14 | 0.23 | 0.10 |
| month:06 | -0.08 | -0.23 | 0.04 | 0.07 |
| month:07 | 0.01 | -0.25 | 0.28 | 0.13 |
| month:08 | 0.10 | -0.04 | 0.27 | 0.08 |
| month:09 | -0.10 | -0.29 | 0.11 | 0.10 |
| month:10 | -0.01 | -0.16 | 0.13 | 0.07 |
| month:11 | 0.01 | -0.09 | 0.12 | 0.05 |
| month:12 | 0.17 | 0.02 | 0.31 | 0.07 |
| Age:Juvenile | 0.08 | -0.12 | 0.24 | 0.09 |
| **Conspecific density** | **-0.30** | **-0.47** | **-0.12** | **0.09** |
| **Conspecific social degree** | **0.22** | **-0.07** | **0.49** | **0.14** |
| **Heterospecific density** | **0.21** | **0.03** | **0.38** | **0.09** |
| **Heterospecific social degree** | **-0.05** | **-0.38** | **0.26** | **0.16** |
